# IP6 and PF74 affect HIV-1 Capsid Stability through Modulation of Hexamer-Hexamer Tilt Angle Preference

**DOI:** 10.1101/2024.03.11.584513

**Authors:** Chris M Garza, Matthew Holcomb, Diogo Santos-Martins, Bruce E. Torbett, Stefano Forli

## Abstract

The HIV-1 capsid is an irregularly shaped complex of about 1200 protein chains containing the viral genome and several viral proteins. Together, these components are the key to unlocking passage into the nucleus, allowing for permanent integration of the viral genome into the host cell genome. Recent interest into the role of the capsid in viral replication has been driven by the approval of the first-in-class drug lenacapavir, which marks the first drug approved to target a non-enzymatic HIV-1 viral protein. In addition to lenacapavir, other small molecules such as the drug-like compound PF74, and the anionic sugar inositolhexakisphosphate (IP6), are known to impact capsid stability, and although this is widely accepted as a therapeutic effect, the mechanisms through which they do so remain unknown. In this study, we employed a systematic atomistic simulation approach to study the impact of molecules bound to hexamers at the central pore (IP6) and the FG-binding site (PF74) on capsid oligomer dynamics, compared to *apo* hexamers and pentamers. We found that neither small molecule had a sizeable impact on the free energy of binding of the interface between neighboring hexamers but that both had impacts on the free energy profiles of performing angular deformations to the pair of oligomers akin to the variations in curvature along the irregular surface of the capsid. The IP6 cofactor, on one hand, stabilizes a pair of neighboring hexamers in their flattest configurations, whereas without IP6, the hexamers prefer a high tilt angle between them. On the other hand, having PF74 bound introduces a strong preference for intermediate tilt angles. These results suggest that structural instability is a natural feature of the HIV-1 capsid which is modulated by molecules bound in either the central pore or the FG-binding site. Such modulators, despite sharing many of the same effects on non-bonded interactions at the various protein-protein interfaces, have decidedly different effects on the flexibility of the complex. This study provides a detailed model of the HIV-1 capsid and its interactions with small molecules, informing structure-based drug design, as well as experimental design and interpretation.

## Introduction

HIV-1 is a structurally complex virus, with not one but two protective shells surrounding the viral RNA genome. While fusion with the host cell membrane is mediated by the outer matrix layer, the inner capsid layer (i.e.: the viral capsid) retains a complex role in viral RNA delivery to the nucleus, depending on several host factors while evading others along the way^(1–3)^. Despite being among the most studied viruses, many questions remain regarding the timely rupture of the viral capsid inside the host cell’s nucleus and the mechanisms through which host factors and therapeutics impact its infectivity.

The mature viral capsid is formed by multiple repeats of the same capsid protein (CA), which contains an N-terminal domain (NTD, residues 1–145) and a C-terminal domain (CTD, residues 150–231), connected by a flexible linker region^(4)^ (Fig. 1D). The viral capsid adopts a variable, asymmetric geometry called a fullerene cone^(5,6)^ (Fig. 1A), composed of around 200 CA hexamers (H) and exactly 12 CA pentamers (P). The irregularity of the fullerene cone is unusual for viral coatings and precludes the use of several experimental techniques used to facilitate detailed studies of such massive multi-protein complexes, posing significant challenges for experimental and computational studies alike. Low-resolution models of empty viral capsids have been published^(7)^, which along with high-resolution structures of the hexamer^(4,8)^ and pentamer subunits^(9,10)^, inform our incomplete understanding of the capsid.

**Fig. 1.**
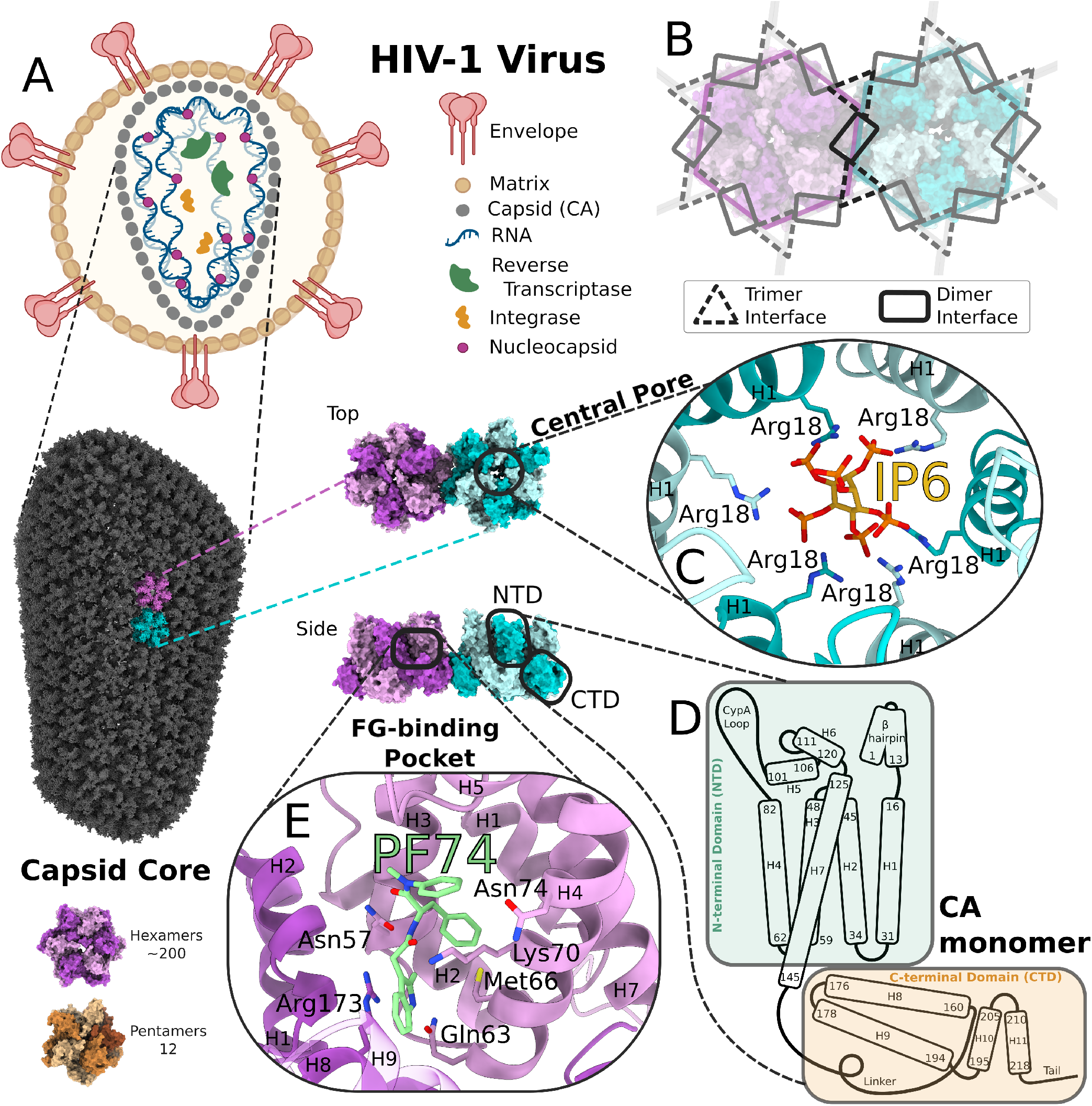
Overview of capsid structural features. A) Viral particle model (created with Biorender.com) with intact capsid cone (PDB:3J3Q). A pair of hexamers (HH) are highlighted in blue and purple for scale. B) The trimer and dimer interfaces, which together form the inter-oligomer interface, are outlined on the HH model. Viewed from above, or from outside the virus, the bundle of NTDs are most visible including the central pore which is shown in greater detail in C) with Arg18 side chains from each capsid chain shown as sticks interacting with the host factor inositolhexakisphosphate (IP6). The structure of the capsid (CA) monomer is shown in D) with helices labeled with “H” and residue numbers for the first and last residue in each secondary structural feature. The FG-binding site is shown in greater detail in E), including key residues shown as sticks and the small molecule PF74 (PDB:4XFZ).

In both hexamer and pentamer oligomers, the collection of 5/6 NTDs form the core of the oligomer with a pore passing through the center (Fig. 1C). A narrow ring of arginines deep within the central pore forms key interactions with cellular components such as inositolhexakisphosphate^(11,12)^ (IP6), FEZ1 ^(13,14)^, and dNTPs^(15,16)^ at various points during the replication process. In the absence of IP6, only unclosed rods of hexamers have been observed to form from a concentrated solution of free capsid proteins^(17–19)^, but the addition of IP6 is sufficient to induce formation of fullerene cones and maintain their stability^(20,21)^. IP6 molecules are packaged with newly forming viruses and are a critical factor in maturation of the virus into its infectious form, including assembly of the capsid fullerene cone^(11,22,23)^. Aside from IP6, it is unknown if other host factors involved in viral replication alter capsid assembly or stability.

Neighboring CA oligomers are attached noncovalently at interfaces formed by contacts between opposing CTDs. The flexible linker connecting CTD and NTD wraps around and forms non-bonded contacts with both the NTDs of other chains in its own oligomer and across to the CTDs of neighboring oligomers. Parts of the CTD and NTD from one chain (helices 2, 7, 8, and 9) wrap around and form a pocket with the NTD of the next chain (helices 3 and 4) (Fig. 1E). This pocket is called the FG-binding pocket due to the capsid’s interaction with host cell proteins Sec24c^(24)^, CPSF6 ^(25,26)^, and Nup153 ^(25,27)^, which all bind through a shared phenylalanine-glycine (FG) motif. The small molecule PF-3450074 (PF74)^(25,28)^ and its more complex derivative lenacapavir^(29,30)^, recently approved for clinical applications, bind to the FG-binding pocket by mimicking the phenylalanine-glycine motif and have been shown to stabilize the capsid and induce early rupture^(31)^. While the effect that PF74 and IP6 have on capsid structural stability hint at the possibility of capsid to be meaningfully impacted by other molecules, the precise mechanism through which either small molecule achieves this effect remains largely speculative due to experimental studies being limited by the size, asymmetry, and relative fragility of the capsid cone *in vitro*. Previous computational studies of the capsid have been challenging for similar reasons. In order to solve the whole-capsid models (PDB: 3J3Q, 3J3Y)^(7)^, Schulten and colleagues performed the landmark all-atom simulations of the entire capsid, which they fit to the low-resolution density maps using MDFF. Due to exorbitant computational cost, simulations of the entire viral capsid^(7,32,33)^ (each >44 million atoms) remain impractical for free-energy calculations. Previous multi-oligomer studies^(34,35)^ have explored thermal stability, ion transport, and non-bonded potential energies at the protein-protein interfaces, however these studies do not account for the potential impact of varying curvatures and approximate entropic effects. However, the majority of all-atom simulation studies have been focused on one CA dimer^(36,37)^, hexamer^(15,38,39)^, or pentamer^(39)^.

A considerable body of work has also been dedicated to modelling the viral capsid at the coarse-grained level. The majority of the coarse-grained capsid models published to date are oriented toward obtaining parameters to replicate assembly of fullerene cones, tubes, and sheets from dimers^(40–47)^. A study of capsid disassembly by Yu et. al^(33)^ simulated the outcome of increasing internal pressure of several intact capsids modeled from cryo-ET densities.

In this work, we seek to bridge the existing gap between allatom studies of single oligomers and coarse-grained models of the entire capsid, by studying both possible pairs of neighboring oligomers (hexamer-hexamer and hexamer-pentamer) at the all-atom level across the range of curvatures found in the experimentally-derived fullerene cone models. The results of this analysis provide unifying perspectives on native viral capsid behavior and the actions of small molecules on coexisting intra- and inter-oligomer interactions, and can inform hypothesis generation for future experimental and computational studies.

## Results

The goal of our MD studies was to characterize every proteinprotein interface found in the intact viral capsid, including their variations induced by different degrees of curvature. Since the minimal system including each of the possible protein-protein interfaces is a pair of neighboring oligomers, we built *apo* and *holo* hexamer-hexamer (HH), and hexamer-pentamer (HP) systems which were interrogated in explicit solvent in the controlled environment of the all-atom simulation. This allowed us to compare the thermodynamic stability of various states and the non-bonded interactions at each protein-protein interface at the atomic scale.

The *apo* hexamer-hexamer model (HH_apo_) was built from a carefully selected pair of hexamer neighbors from the cryoEM capsid tube model (PDB: 3J34 ^(7)^). Although capsid tubes are not closed at the ends, and are therefore not capable of contributing to viral infection, their symmetry allows for resolving higher-quality models than the irregular fullerene cones found in infectious particles, and have therefore been used extensively as a proxy. The *apo* hexamer-pentamer model (HP_apo_), however, could not be built from the same structure given that capsid tubes are formed entirely by hexamers. Therefore, we searched the two available capsid fullerene cone models (PDB: 3J3Q, 3J3Y)^(7)^ to identify hexamer-pentamer pairs with relatively average features by comparison to other hexamer-pentamer pairs. Further details are available in the Supplementary Information. The IP6-bound and PF74-bound hexamer systems (HH_IP6_ and HH_PF74_, respectively) were built using the PDB:6BHS^(20)^ and PDB:4XFZ^(4)^ *holo* hexamer structures, respectively. In each case, two copies of each hexamer structure were fit to the *apo* hexamer-hexamer system based on RMSD alignment of the four chains directly involved at the hexamer-hexamer interface. Due to the lack of any *holo* pentamer models, they were not included in this analysis.

To establish free energies of binding, we first employed steered molecular dynamics (SMD) to separate the oligomers in a dimer along the axis connecting their centers of mass. Then, frames sampled during the SMD simulations were selected to yield evenly spaced windows (1 Å apart) for umbrella sampling (US), with the reaction coordinate starting from the intact pair of oligomers and ending with fully separated oligomers (Fig. 2A). Our studies assume that the hexamer (or pentamer) unit stays intact during disassembly. Similarly, SMD was performed along tilt and twist coordinates, describing the angular differences between the planes of each oligomer (Fig. 2B,C), following conventional definitions^(9)^. The range of HH interface tilt/twist angles we calculated (Supplementary Figure 1) from the whole-capsid models (PDB:3J3Q, 3J3Y)^(7)^ exceed the previously reported range of (-1° to 29°) tilt and (-12° to +12°) twist^(9)^, so we expanded the range of tilt/twist angles evaluated in our simulations to (0° to 45°) and (-15° to +15°), respectively. During this sampling, the other angular degree of freedom was restrained to 0°. The free energy profiles of the two angular coordinates were then summed to yield bidimensional free energy landscapes (Fig. 3A,B), whereupon this landscape could be projected onto a whole-capsid experimental structure (PDB:3J3Q)(Fig. 3C) in order to visualize the local stability of an intact viral capsid. This approximation assumes a lack of coupling between these degrees of freedom, which would be exceedingly computationally expensive to sample, and additionally excludes separation distance, for which no system had a significantly different local minimum. Aside from the free energy landscapes that resulted from these calculations, the trajectories also contain valuable information about the frequency of non-bonded interactions, and we analyze their differences on a per-system basis (Fig. 4), and between tilt angles within a system (Fig. 5). What follows is a detailed analysis of the results for each system considered.

**Fig. 2.**
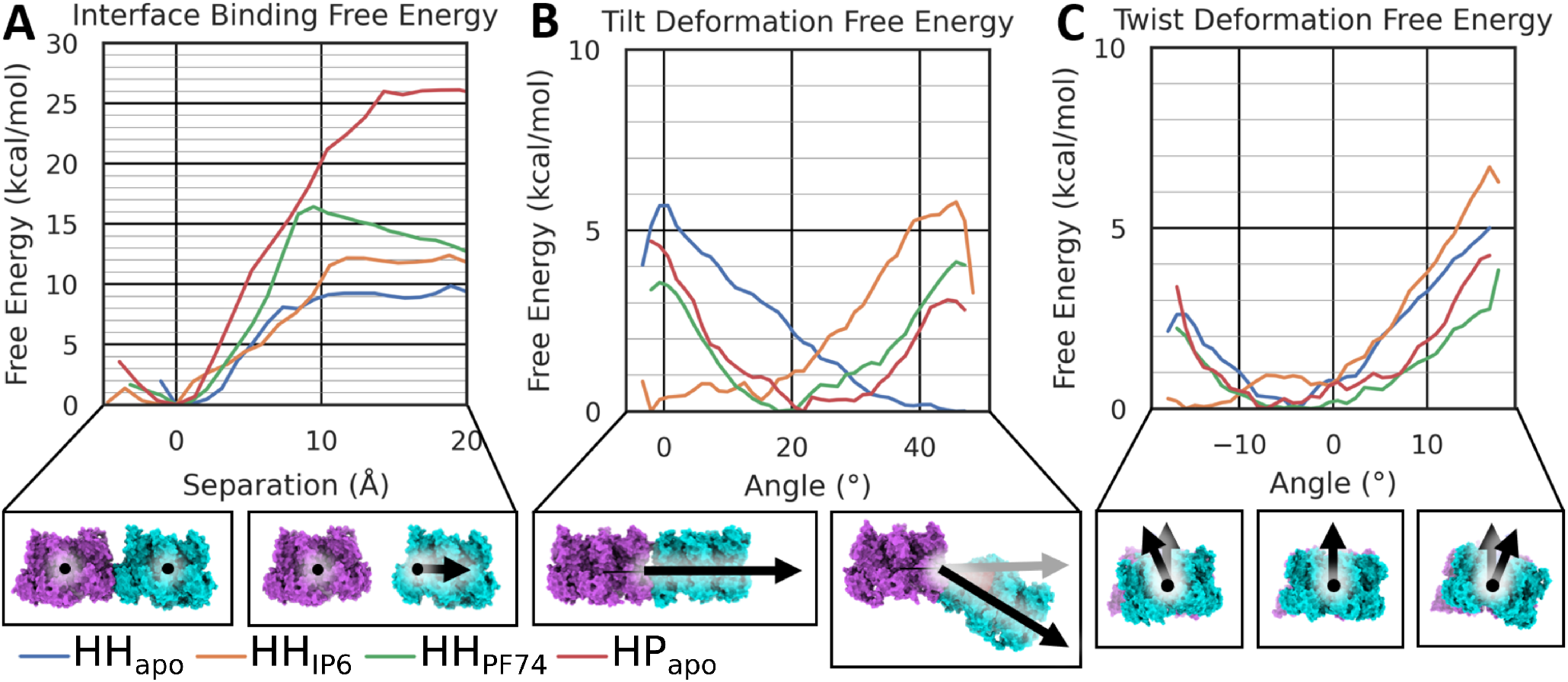
Dissociation and deformation of oligomer pairs. Free energy profiles were calculated from umbrella sampling for four systems: apo hexamer-pentamer (red), and hexamer-hexamer bound to PF74 (green), bound to IP6 (orange), and apo (blue). A) Dissociation free energy calculated over the separation of the centers of mass of each oligomer; Free energy of deformation as a function of tilt B) and C) twist angle.

**Fig. 3.**
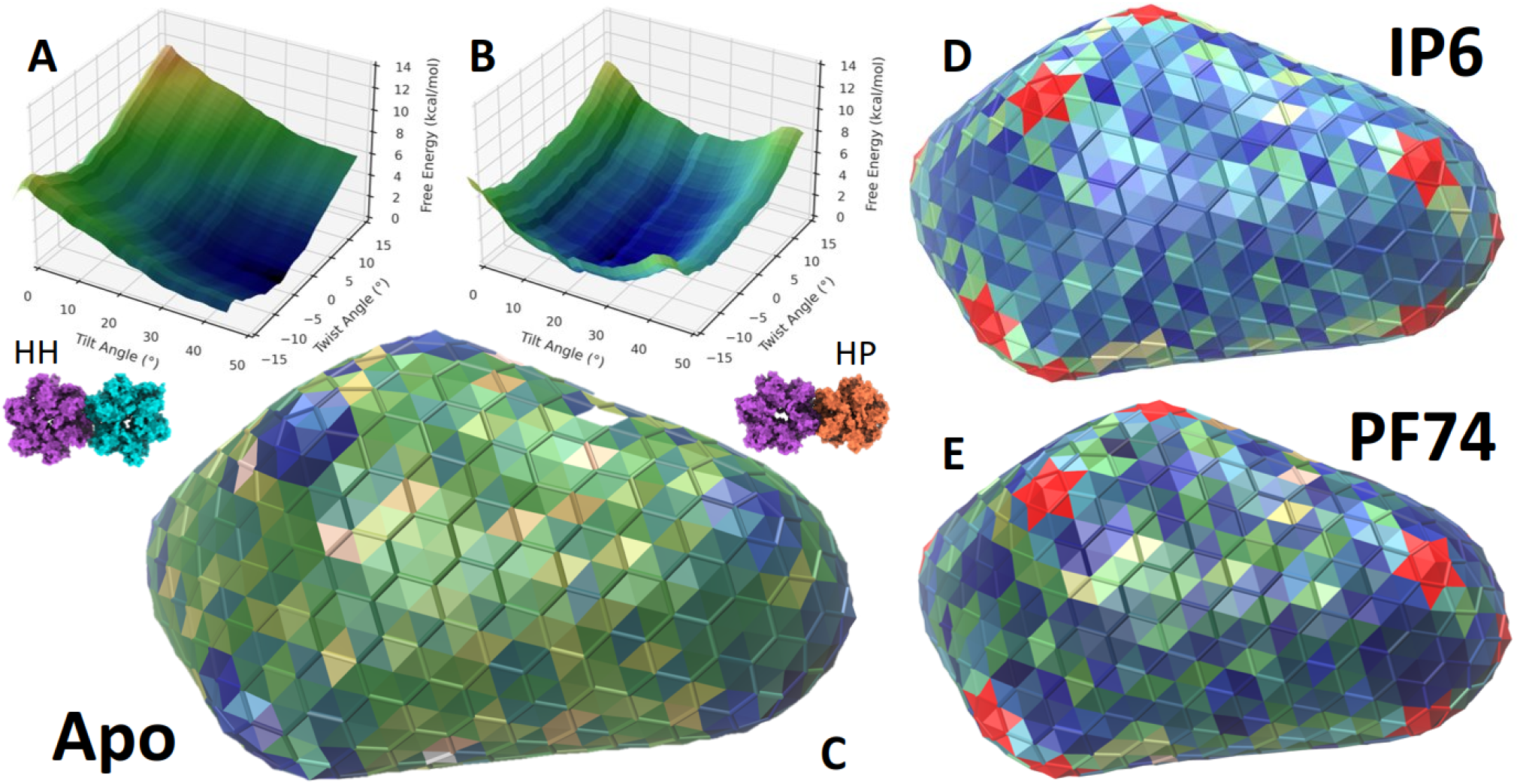
Free Energy of each oligomer pair introduced by tilt/twist curvature, as depicted on a full capsid model. Tilt/Twist Free Energy landcapes for A) HH_apo_ and B) HP_apo_ oligomer-of-oligomers, by summation of each degree of freedom. Landscapes for C) HH_apo_ and HP_apo_, D) HH_IP6_, and E) HH_PF74_ are plotted onto a wirecage model derived from the (PDB:3J3Q) whole capsid model. *holo*-HP interfaces are in red where free energy data is missing.

**Fig. 4.**
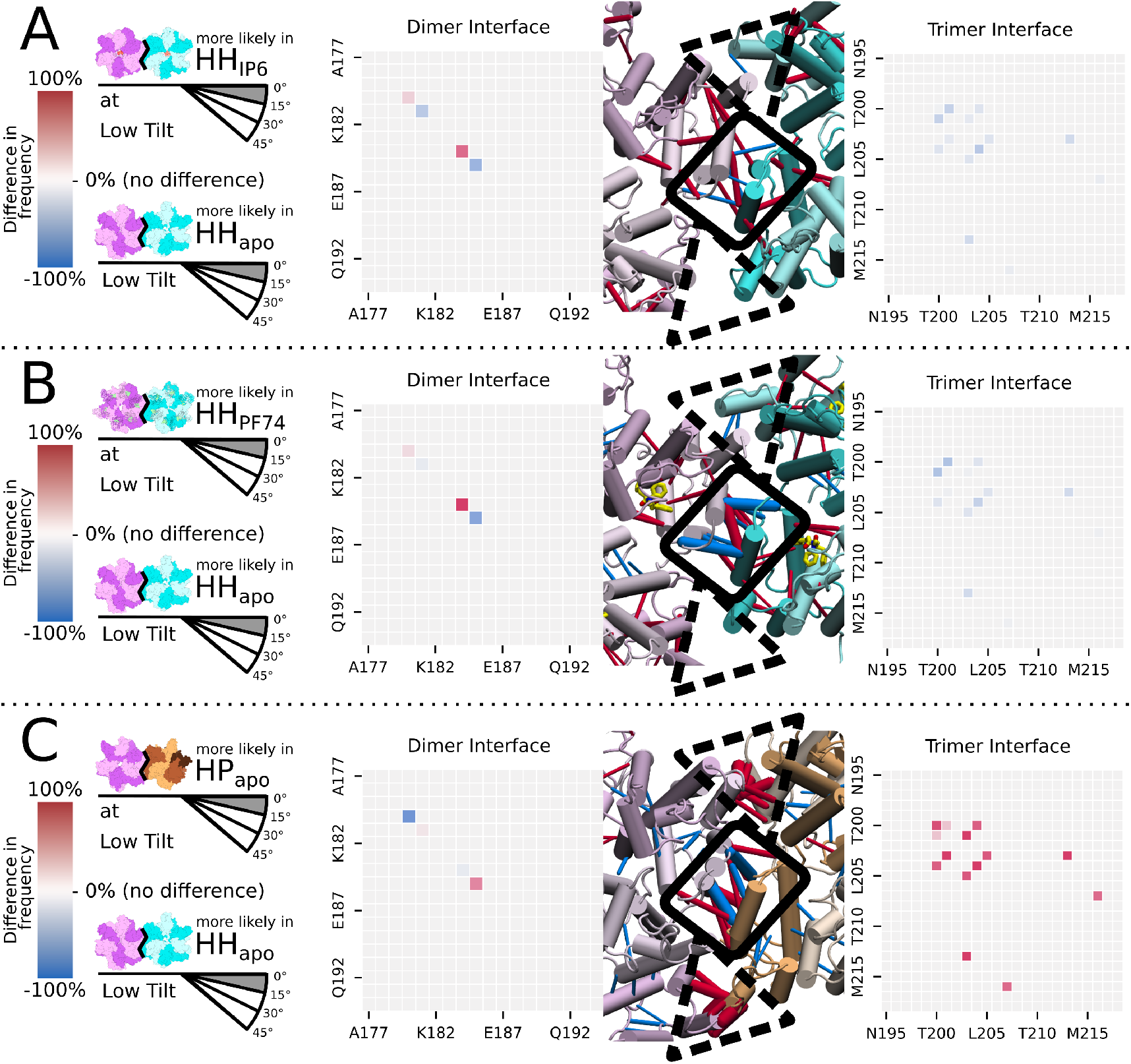
Difference in frequency of non-bonded interactions present at interface of HH or HP (inter-oligomer) systems at low tilt angles. Values fall into the range from +100% (red) to -100% (blue) where zero indicates there is no difference between the systems, but not necessarily that the interaction was absent. A,B,C) Relative interaction frequency for HH_IP6_, HH_PF74_, and HP_apo_, with respect to HH_apo_, using data from frames falling into the low tilt (0-15°) range. Left panel: heatmap of differences associated with helix 9, representative of the dimer interface, in order of the amino acid sequence. Right panel: heatmap of differences at the trimer interface, including helices 10 and 11. Middle panel: structure of the inter-oligomer interface with hexamer (lavender and cyan) and pentamer (beige) helices in cylinders. Dashed black triangles indicate the trimer interface and solid black boxes indicate the dimer interface. More (red) and less (blue) frequent interactions are shown as rods whose diameter reflects the average interaction frequency.

**Fig. 5.**
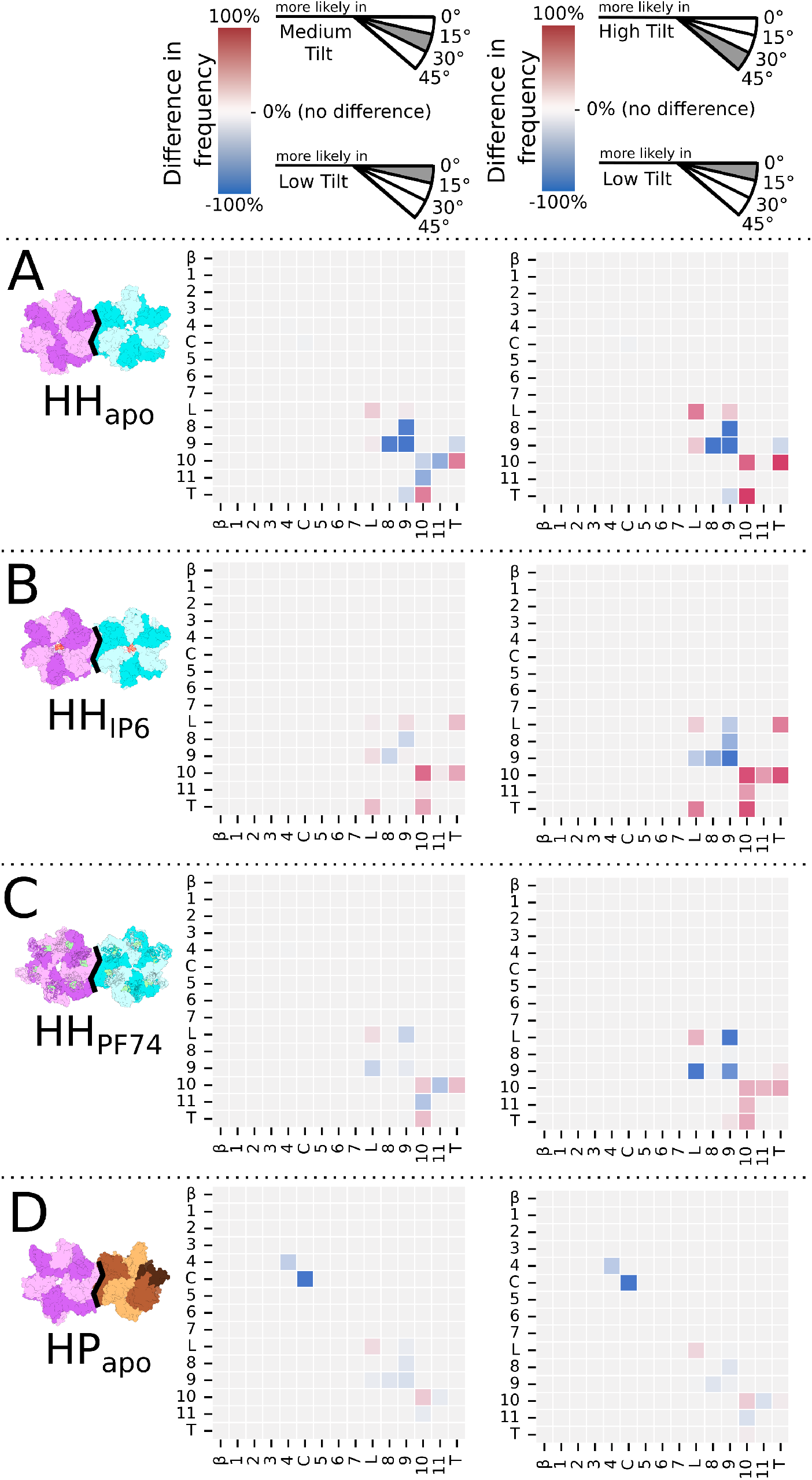
Difference in frequency of non-bonded interactions between secondary structural features at interface of HH or HP (inter-oligomer) systems over increasing tilt angles. Values fall into the range from +100% (red) to -100% (blue) where zero indicates there is no difference between the systems, but not necessarily that the interaction was absent. A-C) Differences between a system in low tilt (0-15°) and either medium (15-30°, left column) or high tilt (30-45°, right column). Values are determined by the collective difference in frequencies of each of the residues in a segment as defined by Fig. 1D.

### HP interface is more stable than HH interface

Our results suggest that the HP interface is considerably stronger (up to three-fold) than the HH interface. The HP interface required 25 kcal/mol to separate hexamer from pentamer, whereas breaking two hexamers apart required only 9 kcal/mol (Fig. 2A). Intact HP_apo_ interfaces bear a vastly different network of non-bonded interactions than any of the HH systems. The trimer interface in the HP_apo_ system was by far the most well-connected of any system we studied (Fig. 4D). There were a mixture of differences between HH_apo_ and HP_apo_ at the dimer interface, both in increased and decreased interaction frequencies, for residues involving both helix 9 and the linkers (see Fig. 4D and Supplementary Figure 3,7,8).

### HP pairs are optimally bent in intact viral capsid

The HH_apo_ system was found to have a significant preference for higher tilt angles, with the maximum of 5.6 kcal/mol at 0° and effectively minimum values higher than 30° (Fig. 2B). The HP_apo_ system, on the other hand, disfavored both high (+3 kcal/mol) and low (+4.5 kcal/mol) tilt extremes, with its minimum at 20° tilt (Fig. 2B). The free energy across twist angles revealed a similar asymmetric curve in both HH_apo_ and HP_apo_ systems with a minimum at -5° twist and a maximum at +15° twist (+5 kcal/mol) Fig. 2C.

Interestingly, the HP_apo_ system formed non-bonded contacts between the NTDs across the HP interface involving helix 4 and the CypA loop (Fig. 5E). This is indicative of crowding between the NTDs between adjacent oligomers, which are typically assumed to not be involved in inter-oligomer interactions.

In our simulations, the ideal angles measured for the HP_apo_ system closely match their distribution in the whole viral capsid models, with tilt and twist minima overlapping the naturally occurring interface angles almost exactly (Supplementary Figure 1). In fact, very few of the HP interfaces found in the currently available viral capsid models^(7)^ would deviate more than 1 kcal/mol from the ideal minima we identified (Fig. 3B,C). When mapping the energetic values on one of the *apo* capsid models (Fig. 3C), they appear as islands of stability among the otherwise mostly unstable HH interfaces.

### IP6 stabilizes flat patches

Overall, our simulation results suggest that HH_apo_ is intolerant to flattening. Indeed, when mapping the calculated HH_apo_ tilt/twist free energy profile values on the viral capsid models, it appears that large swaths of hexamers across the midsection appear to be several kcal/mol away from their theoretical energetic minima (Fig. 3A,C).

However, the addition of IP6 dramatically changes the energetic landscape throughout the viral capsid, converting the flatter sections to low-energy states (Fig. 3D). This is a result of an inversion of the free energy profile over tilt angles, with favorable states from 0° to 20° and a steadily rising barrier up to 5.3 kcal/mol at 40° (Fig. 2B).

When binding, IP6 establishes direct contacts with the *β*hairpin and helix 1, providing stability to the NTDs (Supplementary Figure 6). Yet, the effect of IP6 goes beyond the NTD, increasing the frequency of key contacts between helix 3 and helix 8 (residues Asn57, Val59 to Arg173) from a neighboring chain (Supplementary Figure 6). In turn, the contacts throughout the CTD were increased, especially between helix 8, 9, and 11 (Supplementary Figure 9). Ultimately, helix 9 and the linker region establish a mixture of more prevalent interactions across the dimer interface to the neighboring hexamer (Fig. 4B), including the hydrophobic interaction between the two opposing Trp184’s, and less prevalent interactions, including the hydrophobic interaction between the two opposing Met185 side chains. Finally, at the inter-oligomer trimer interface, which includes contacts between opposing helices 10 and 11 (Fig. 4B), the HH_IP6_ system showed several mildly decreased non-bonded interactions.

Our simulations showed that compared to HH_apo_, the HH_IP6_ dimer interface better withstood loss of non-bonded interactions by deformation to high tilt angles, especially at the helix 8 helix 9 interface (Fig. 5C). Additionally, HH_IP6_ gained more non-bonded contacts at the trimer interface over both medium and high tilt angles than HH_apo_, including contacts between helices 10 and 11, which are not seen in HH_apo_ (Fig. 5C).

### PF74 rigidifies hexamer-hexamer interactions

The tilt free energy profile of HH_PF74_ more closely matches the free energy landscape of the HP_apo_ system than the HH_apo_ system (Fig. 2B). With a minima around 20° tilt, PF74-bound HH pairs face barriers to both flattening and further bending Fig. 2B. When analyzing intact viral capsids (PDB:3J3Q)^(7)^ in light of the free energy profile of HH_PF74_, our data predicts that binding of PF74 alone to the viral capsid would introduce an irregular pattern of stability and instability across the midsection of the capsid cone, and further stabilize the curved end caps, most notably the narrow end (Fig. 3E).

The interactions between residues Arg173 and Asn57 and Val59, which form the floor of the PF74 binding site, appear more frequently in the HH_apo_ system, and to a similar degree as in the HH_IP6_ system (Supplementary Figure 6). Similarly, non-bonded interactions throughout the CTD increased, with major contributions from helices 8, 9, and 11 (Supplementary Figure 9). At low tilt angles, the profile of non-bonded interactions at the HH_PF74_ inter-oligomer interface was strikingly similar to HH_IP6_ as compared to HH_apo_, but HH_PF74_ experienced greater losses at the dimer interface and less gains at the trimer interface with increasing tilt angle. However, when considering the entire data set (Supplementary Figure 3), HH_PF74_ and HH_IP6_ adopt very similar non-bonded interactions profiles at the inter-oligomer interface.

## Discussion

The mature viral capsid is a complex, asymmetric and often irregular structure exhibiting varying degrees of curvature, despite being composed of a single protein (capsid protein, CA). This protein self-assembles in 5- and 6-membered oligomers, and the interface between them is found in a wide range of conformations. Consequently, we hypothesized that some oligomer interfaces in the mature capsid are more stable than others.

The most conspicuous loci of structural variation are the pentamers, and indeed we found that the HP interface was considerably more resistant to complete separation than the HH interface, and that pentamers naturally occur in ideal tilt and twist angles for HP interfaces in the fullerene cone geometry. In contrast, the hexameric part of the capsid is dotted with regions of relative instability, owing to tilt and twist deviations from ideality. However, the addition of either IP6 (in the central pore) or PF74 (at the FG-binding site) increases the overall stability through modulation of the tilt/twist angle preferences. In HH_IP6_, we observed that IP6 inverted the free energy profile with respect to the tilt angle of the HH_apo_ complex, making low tilt angles more favorable. Therefore, one of the roles IP6 may play in the viral life cycle is to stabilize the otherwise energetically unfavorable flat portions of the viral capsid, allowing for the coexistence of curved low-energy states (*apo*) and flat low-energy states (with IP6 bound). This is supported by prior evidence that viral capsids are more stable when incubated in solution containing IP6 ^(12,22,48)^, and that formation of new closed capsids *in vitro* is only possible in the presence of IP6 ^(20)^. Indeed, without IP6 only cylindrical rods composed entirely of hexamers can be made *de novo* from capsid subunits in solution^(5,17–19,21)^. This is consistent with the tilt preferences we observed for HH_apo_ and HH_IP6_ interfaces. Together with the fact that small molecules affected tilt preferences more than dissociation or twist free energies, this leads us to postulate that tilt angle modulation is the primary mechanism by which IP6 and PF74 affect capsid stability.

IP6 may also play a role in the timely disassembly of the viral capsid. In order for infection to occur, the RNA inside the viral capsid must be reverse-transcribed into DNA, and subsequently the viral capsid must disassemble, allowing the nascent DNA to be integrated into the host cell’s genome. Previous studies have found that reverse transcription occurs while the capsid is still intact^(12,16,49)^ and that the process leads directly to capsid disassembly^(12,50)^.

In order for this to be possible, individual dNTPs from the host cell cross through the capsid barrier^(16)^ to supply the substrate for DNA synthesis. In one previously proposed mechanism, dNTPs pass through the central pore by first binding to Arg18 alongside IP6, then turn around while squeezing through the ring of Arg18 side chains to bind a ring of Arg25 side chains, before being released into the capsid interior^(15)^. Given that the full viral genome requires the formation of 8.8 kbp of dsDNA, some 8,800 dNTPs must translocate into the viral capsid through the central pores of the roughly 200 hexamers. Given the mutually repulsive negative charges of IP6 and dNTP, the remodelling of the NTDs needed to accommodate the dNTP^(15)^, and the disturbance of favorable interactions between IP6 and Arg18, it seems reasonable to suggest that some IP6 molecules could be displaced from their hexamers.

At the same time, the growing dsDNA strand applies greater outward pressure on the capsid walls than the previous single stranded RNA, which together with increasing, irregular capsid instability provides ample opportunity for cracks to form, leading to capsid rupture.

While addition of PF74 to HH oligomers increases the overall stability of the capsid, the range of favorable tilt angles becomes relatively narrow around 20°, and deviations from the favorable range face a steeper energy barrier than for any of the other tilt energy landscapes. Therefore, as a result of the reduced angular tolerance, PF74 makes the HH interface more rigid and brittle. This rigidification is in line previous studies which have shown that PF74 accelerates formation of openings in the capsid wall while simultaneously stiffening intact sections^(31,51)^.

Another mechanism of action of PF74 is inhibition of formation of mature capsid from infected cells^(28)^. Our results regarding the tilt preferences of HH systems offer a plausible mechanism for this effect. The narrow preference for tilt angles around 20°is concomitant with energetically unfavorable flat (0°) geometries, reducing the likelihood of formation of the flat hexameric patches in mature capsid where numerous HH neighbors need a tilt angle around 0°.

The correlation between interaction patterns and modulation of tilt angles by small molecules is not immediately obvious. On one hand, when comparing HH_PF74_ to HP_apo_ we observed similar tilt preferences but distinct interaction profiles. However, we observed the opposite when comparing the effect of IP6 to that of PF74 on HH pairs, with different tilt preferences emerging from similar interaction profiles. The changes in interactions induced by PF74 and IP6 are strikingly similar, despite distal binding sites, and both small molecules make medium to low tilt angles more favorable, thereby stabilizing flat patches of the capsid. This includes the previously studied^(52)^ Arg173 to Asn57 and Val59 pair of hydrogen bonds (Supplementary Figure 5) which form a bridge from the NTD core to the CTDs and inter- and intra-oligomer interactions between Trp184 and Met185 (Fig. 4A,B), previously used as a key motif for small molecule inhibitor design^(53)^. Therefore, we hypothesize that the change in interactions induced by IP6 or PF74 may serve as an *in silico* marker for tilt angle modulation, which in turn is likely to affect capsid stability. This may aid in future drug discovery campaigns seeking to expand off the first-in-class success of lenacapavir, and provide useful descriptors to predict the phenotypic effect of drug candidates. This is especially true as these interactions should be identifiable even from relatively short, unbiased simulations in presence of small molecules. In turn, this opens up the possibility of virtual screening on this target by highlighting regions of high interaction frequency variability that would be both more susceptible to modulation, and more effective in perturbing the overall viral capsid behavior.

In summary, this work uses MD simulations to investigate the mechanism of action of a small molecule host factor and a drug-like compound with clinical implications for HIV-1 infection. The determinants of their action on such a nonenzymatic target has been challenging to disentangle due to intrinsic difficulties associated with the complex and dynamic arrangement of the oligomeric states of the HIV-1 capsid protein, the sole building block responsible of the viral capsid properties. Our approach allowed us to identify a previously unreported effect of small molecules on the angular preference between capsid oligomers when assembled to form the whole viral capsid.

Together with the analysis of residue interactions at oligomer interfaces, the revealed susceptibility of tilt angle preferences to small molecules will help the formulation of new hypotheses and interpretation of results of future studies of the HIV-1 capsid.

## Methods

### System Preparation

#### Structure

For each system to be simulated, different experimental structures were selected and manipulated to assemble the starting coordinates of the simulations.

#### HH_apo_

Chains A-L were taken from PDB: 3J34 ^(7)^ to build the apo two-hexamer (HH) system. They were rotated and translated in Cartesian space such that the center of mass of one hexamer (chains A-F) is located at the origin and the center of mass of the second hexamer (chains G-L) lies on the X axis in VMD^(54)^.

#### HP_apo_

The hexamer-pentamer (HP) system was selected from whole capsid model in PDB: 3J3Q^(7)^. Of the 120 HP interfaces found in the 3J3Y^(7)^ and 3J3Q^(7)^ whole capsid structures, one HP pair was chosen through a combination of its interface tilt/twist angles as well as the presence of its nonbonded interactions (see Supplemental Information).

*Holo*-pentamer structures were not included in this study due to unavailability of structures and recent findings^(10,55)^ in which pentamers were not found to bind any substituents.

#### HH_IP6_

The PDB: 6BHS^(20)^ was used to build the HH system with IP6 bound. This structure was manually mutated in several positions to restore wild type residues (see Supplementary Information) and at A92E and G208A to match the sequence of the apo structure (PDB: 3J34). Missing loops 88-89 and 177-186 were rebuilt by RMSD best-fit alignment to the adjoining residues in the HH_apo_ model. The C-terminal tail from residue 219 to 231 was also rebuilt. The HH system was built by aligning two copies of the *holo*-hexamer onto the HH_apo_ system using the backbone of the four chains directly involved in the interface as a template. Chains were relabeled to match the scheme of the HH_apo_ system. We included one IP6 molecule for simplicity and based on structural studies indicating that only one IP6 molecule (in the higher position above Arg18) is common in capsids formed inside native virions^(38)^.

#### HH_PF74_

The PF74-containing system was built using the PDB: 4XFZ^(4)^ with only one necessary mutation at G208A to match the sequence of the HH_apo_ system. In the same fashion as the HH_IP6_ system, two copies of the PF74-containing *holo*hexamer were aligned with the HH_apo_ system via RMSD best-fit alignment according to the four chains involved in the HH interface. All 12 PF74 binding sites in the HH complex contained a ligand molecule. Missing loops at 4-12, 222-231 were copied from the HH_apo_ system via RMSD alignment. Chains were relabeled to match the HH_apo_ system.

### Histidine Protonation

The initial protonation states were determined using the PROPKA^(56)^ online server and modified in selected cases for consistency (see Supplemental Information).

### Parameters

Connectivity for the resulting protein structure was produced by the PSFGEN package in VMD^(54)^ using CHARMM36m^(57)^ parameters for the protein, CHARMM36 ^(58)^ for ions, and TIP3P^(59)^ for water. A disulfide bridge was added between Cys198 and Cys218 in each chain. IP6 was protonated at each ring carbon, leaving the phosphates deprotonated for a charge of -12. See Supplemental Information 10,11 for protonated IP6 and PF74 molecules. Parameters for IP6 and PF74 were generated using the CGENFF^(60)^ server with penalties within acceptable values.

### Solvation & Ionization

Using VMD’s solvate tool, 15 Å water was added in all directions beyond the extent of the protein except in the +X dimension which is solvated with 40 Å water to allow for space for separation without creating non-bonded interactions with the next periodic image. Randomly selected waters were then replaced with sodium and chloride ions using VMD’s Autoionize tool to counterionize the protein and bring the total salt concentration to 0.15 M.

### Minimization & Equilibration

All simulations were performed using NAMD 2.12 (CUDA version)^(61,62)^. All systems were minimized for 10,000 steps via steepest descent, followed by a 6-step equilibration totaling to 100 ns (see Supplemental Information). A Langevin thermostat^(63)^ and an isotropic Nosé-Hoover Langevin barostat^(64,65)^ were used, set to 300 K and 1.01325 bar (1 atm), respectively. The barostat oscillation period was 50 fs and the damping time was 25 fs. After the first three steps of equilibration, all simulations were run with a timestep of 2 fs/s with SHAKE applied to all hydrogens. Particle Mesh Ewald^(66)^ was used to calculate long-range electrostatics with a cutoff of 12 Å, a pair list distance of 16 Å, and a switching distance of 10 Å.

### Interface Breaking

#### Definition of Reaction Coordinate

In order to sample over the process of breaking the HH or HP interface, the simplest possible reaction coordinate was defined as the distance between the center of mass of one entire hexamer to the center of mass of its hexamer or pentamer neighbor. Thus, the beginning of the reaction coordinate is not zero but rather 95 Å which represents the distance between the two groups’ centers of mass when the interface is fully intact.

#### Generation of Starting Structures

As described previously, the HH or HP pair were aligned on the x-axis with the center of mass of one group at the origin and the other at some point in the +x direction in VMD. The group at the origin was selectively restrained (see Supplementary Table 3) in order to prevent drift and the second group was pulled along the +x axis via steered molecular dynamics (SMD). The SMD dummy atom was attached to the center of mass of the second group’s backbone with a spring constant of 7 kcal/mol/Å^2^ and was pulled at a constant velocity of 1 Å/ns for 40 ns.

#### Umbrella Sampling

The resulting simulations were used to select representative starting frames for each desired US window, spaced 1 Å apart using a custom script in VMD. Umbrella windows were run for 16 ns each with a simple distance harmonic restraint of 2.5 kcal/mol/Å^2^ and a width of 1 Å. Collective variable values were output every 20 steps or 40 fs.

### Tilt/Twist Deformations

#### Definition of Reaction Coordinates

The simulations which calculate the free energy of performing angular deformations on the HH or HP pair vary over bending and twisting motions with the interface between them intact. In each case, the reaction coordinate is along various angles (in degrees). In order to apply the general definition^(9)^ to the atomistic models herein, we first calculated center of mass of each of the 5 or 6 protein chains, then took the singular value decomposition of the matrix composed of their coordinates in Cartesian space. The resulting right singular vector is the normal vector to the best-fit plane for the hexamer or pentamer.

Then when the HH or HP pair is re-aligned on the x-axis, the “tilt” value is determined by the angle produced by rotation around the z-axis and the “twist” value is determined by the angle produced by rotation around the x-axis. Rotations around the y-axis in this orientation would require a break in the lattice and are therefore not previously reported nor biased in our studies. See Fig. 2 for a visual representation of these reaction coordinates.

Rather than using SMD, which is limited in scope to directional pulls along Cartesian axes, the biased simulations herein used the difference of the two groups’ spin angle collective variables with reference to the z-axis for “tilt” and xaxis for “twist” reaction coordinates.

#### Generation of Starting Structures

In order to robustly define tilt and twist angles during umbrella sampling so that the collective variables reflect the angular deformations of the two groups with reference to each other, the initial apo structure (PDB: 3J34 chains A-J) was biased from its native 9.3° tilt, 9.3° twist to produce a flat structure by biasing individual spin angle restraints by +4.8° tilt, –2.8° twist for hexamer 1 (chains A-F) and –4.5° tilt, +6.5° twist for hexamer 2 (chains G-L) over the course of 400 ps. The frame from the resulting simulations closest to flat (0° tilt, 0° twist) according to geometric post-processing was chosen as the reference for further simulations. Then, three separate simulations used the flat reference structure with the same collective variables to explore the configurations produced by deformations up to +35° tilt, –17° twist, or +17° twist. Biased tilt simulations were run for 40 ns and each biased twist simulation was run for 17 ns. During these simulations, a simple distanceZ restraint along the Y axis from each hexamer’s center of mass to the origin prevents the complex from drifting in the Y direction. Importantly, each of the restraints are orthogonal, allowing for separate decomposition of the sampling. The force constant for Y axis distances was 1 kcal/mol/Å^2^ and for the tilt and twist angles was 2 kcal/mol/degree^2^. HH_IP6_, HH_PF74_, and HP_apo_ systems were prepared in the same fashion with minor differences in starting curvatures.

#### Umbrella Sampling

The resulting simulations were used to select representative starting frames for each desired US window, spaced 1° apart using a custom script. Umbrella windows were run for 16 ns each with a harmonic restraint of 2.5 kcal/mol/degree^2^ and a width of 1°. The Y axis distance restraint is not included in US window restraints but varies minimally.

### Analysis

#### WHAM Analysis

Several custom scripts assist in postprocessing outputs from the collective variables for each window, creating lists of windows, and running the weighted histogram analysis method (WHAM)^(67)^ to generate potentials of mean force (PMFs). Increments of 1 ns were removed from the beginning and/or end of each window to assess equilibration and convergence. A consensus of 6 ns equilibration time was sufficient to reduce changes in the resulting PMF (per ns removed) to less than 1 kcal/mol in all systems and was therefore excluded from each window for further analysis. The same convergence criteria was used to assess total run time for each window, and 16 ns of sampling was found to be sufficient to reduce further changes (per ns added) to the PMF to less than 1 kcal/mol. Thus, the total sampling for each window is 10 ns. Accounting for varying lengths of reaction coordinates for distance, tilt, and twist systems, the total sampling that is directly included in the PMF for HH_apo_, HP_apo_, HH_IP6_, and HH_PF74_ is 4.44 microseconds.

#### Non-bonded Interactions Analysis

The ProLIF^(68)^ package along with MDAnalysis^(69)^ were called in python to detect the presence of hydrogen bonds, salt bridges, pi-pi stacks, cation-pi interactions, and hydrophobic interactions between various protein-protein interfaces in simulation contents saved every 200 ps. From the raw data, we were able to match results of each frame to the specific location along the reaction coordinate that the system was at at that point in time, allowing us to analyze the resulting frequencies with reference to the reaction coordinate.

One reaction coordinate we were especially interested in studying was those assessing various tilt angles. In order to separately evaluate the differences between systems (HH_apo_, HP_apo_, HH_IP6_, HH_PF74_) and over different tilt angles within the same system, we calculated two sets of deltas by averaging the frequencies over the range of 1° bins and then subtracting. The first is the difference in frequency of having found a particular interaction intact between either the HH_IP6_ tilt, HH_PF74_ tilt, or HP_apo_ tilt system and the HH_apo_ tilt system, which serves as our reference, over tilt angles of 0-15° (low tilt). The second set of deltas is by comparison of either the mid tilt (15-30°) or high tilt (30-45°) ranges versus the low tilt (0-15°) range of the same system. In this way, we are able to compare which non-bonded interactions may be relevant to stabilizing both the oligomeric unit (H/P) and the interface between oligomers under different conditions.

The HH or HP interface contains three unique sections: the dimer interface composed of contacts between helix 9 and the linker region of the monomer forming the largest surface of the interface with helix 9 and the linker region of the monomer across; and two asymmetric aspects of the trimer interface formed by the convergence of helix 10 and 11 from three adjoining oligomers (Supplementary Figure 2). Because of the vastly increased simulation time that would have been required to include a third oligomer, and thus be able to evaluate a complete trimer interface, we are unable to make strong conclusions about interactions involved in the trimer interface. That being said, our analysis includes interactions involved in one entire dimer interface plus two incomplete trimer interfaces (together the inter-oligomer interface), as well as 12(H)/11(P) copies of interactions between monomers within each oligomer (intra-oligomer), and between segments of each monomer (intra-monomer), as they are reported in detail in the Supplemental Information.

## Availability

The code used for non-bonded interactions analysis in this manuscript is available at the lab GitHub: https://github.com/forlilab/NBKit.

## ACKNOWLEDGEMENTS

We thank Art Olson, David Goodsell and members of the HIVE Center for insightful discussions. This work was supported by NIH HIVE and B-HIVE Center Grants (AI150472, AI170855) and the University of California San Diego Medical Scientist Training Program T32GM007198 Grant. This is manuscript #xxxxx from Scripps Research.

## AUTHOR CONTRIBUTIONS

C.M.G. performed molecular dynamics simulations and analysis thereof, supervised by B.E.T. and S.F. Paper writing was done by C.M.G., M.H., D.S-M., and S.F.

## Bibliography

1. Zhuang, S.; Torbett, B. E. Interactions of HIV-1 Capsid with Host Factors and Their Implications for Developing Novel Therapeutics. Viruses 2021, 13, 417.

2. Rossi, E.; Meuser, M. E.; Cunanan, C. J.; Cocklin, S. Structure, Function, and Interactions of the HIV-1 Capsid Protein. Life 2021, 11, 100.

3. Saito, A.; Yamashita, M. HIV-1 capsid variability: viral exploitation and evasion of capsid-binding molecules. Retrovirology 2021, 18, 32.

4. Gres, A. T.; Kirby, K. A.; KewalRamani, V. N.; Tanner, J. J.; Pornillos, O.; Sarafianos, S. G. X-ray crystal structures of native HIV-1 capsid protein reveal conformational variability. Science 2015, 349, 99–103, Publisher: American Association for the Advancement of Science.

5. Li, S.; Hill, C. P.; Sundquist, W. I.; Finch, J. T. Image reconstructions of helical assemblies of the HIV-1 CA protein. Nature 2000, 407, 409–13.

6. Ganser, B. K.; Li, S.; Klishko, V. Y.; Finch, J. T.; Sundquist, W. I. Assembly and Analysis of Conical Models for the HIV-1 Core. Science 1999, 283, 80–83.

7. Zhao, G.; Perilla, J. R.; Yufenyuy, E. L.; Meng, X.; Chen, B.; Ning, J.; Ahn, J.; Gronenborn, A. M.; Schulten, K.; Aiken, C.; Zhang, P. Mature HIV-1 capsid structure by cryo-electron microscopy and all-atom molecular dynamics. Nature 2013, 497, 643–6.

8. Pornillos, O.; Ganser-Pornillos, B. K.; Kelly, B. N.; Hua, Y.; Whitby, F. G.; Stout, C. D.; Sundquist, W. I.; Hill, C. P.; Yeager, M. X-ray Structures of the Hexameric Building Block of the HIV Capsid. Cell 2009, 137, 1282–1292.

9. Mattei, S.; Glass, B.; Hagen, W. J. H.; Kräusslich, H.-G.; Briggs, J. A. G. The structure and flexibility of conical HIV-1 capsids determined within intact virions. Science 2016, 354, 1434–1437, Publisher: American Association for the Advancement of Science.

10. Stacey, J. C. V.; Tan, A.; Lu, J. M.; James, L. C.; Dick, R. A.; Briggs, J. A. G. Two structural switches in HIV-1 capsid regulate capsid curvature and host factor binding. Proceedings of the National Academy of Sciences 2023, 120, e2220557120.

11. Renner, N.; Mallery, D. L.; Faysal, K. M. R.; Peng, W.; Jacques, D. A.; Böcking, T.; James, L. C. A lysine ring in HIV capsid pores coordinates IP6 to drive mature capsid assembly. PLOS Pathogens 2021, 17, e1009164.

12. Mallery, D. L.; Márquez, C. L.; McEwan, W. A.; Dickson, C. F.; Jacques, D. A.; Ananda-padamanaban, M.; Bichel, K.; Towers, G. J.; Saiardi, A.; Böcking, T.; James, L. C. IP6 is an HIV pocket factor that prevents capsid collapse and promotes DNA synthesis. eLife 2018, 7, e35335, Publisher: eLife Sciences Publications, Ltd.

13. Huang, P.-T.; Summers, B. J.; Xu, C.; Perilla, J. R.; Malikov, V.; Naghavi, M. H.; Xiong, Y. FEZ1 Is Recruited to a Conserved Cofactor Site on Capsid to Promote HIV-1 Trafficking. Cell Reports 2019, 28, 2373–2385.e7.

14. Malikov, V.; da Silva, E. S.; Jovasevic, V.; Bennett, G.; de Souza Aranha Vieira, D. A.; Schulte, B.; Diaz-Griffero, F.; Walsh, D.; Naghavi, M. H. HIV-1 capsids bind and exploit the kinesin-1 adaptor FEZ1 for inward movement to the nucleus. Nature Communications 2015, 6, 6660, Number: 1 Publisher: Nature Publishing Group.

15. Xu, C.; Fischer, D. K.; Rankovic, S.; Li, W.; Dick, R. A.; Runge, B.; Zadorozhnyi, R.; Ahn, J.; Aiken, C.; Polenova, T.; Engelman, A. N.; Ambrose, Z.; Rousso, I.; Perilla, J. R. Permeability of the HIV-1 capsid to metabolites modulates viral DNA synthesis. PLoS Biol 2020, 18, e3001015.

16. Jacques, D. A.; McEwan, W. A.; Hilditch, L.; Price, A. J.; Towers, G. J.; James, L. C. HIV-1 uses dynamic capsid pores to import nucleotides and fuel encapsidated DNA synthesis. Nature 2016, 536, 349–53.

17. Ganser-Pornillos, B. K.; Von Schwedler, U. K.; Stray, K. M.; Aiken, C.; Sundquist, W. I. Assembly Properties of the Human Immunodeficiency Virus Type 1 CA Protein. Journal of Virology 2004, 78, 2545–2552.

18. Gross, I.; Hohenberg, H.; Kräusslich, H.-G. In Vitro Assembly Properties of Purified Bacterially Expressed Capsid Proteins of Human Immunodeficiency Virus. European Journal of Biochemistry 1997, 249, 592–600.

19. von Schwedler, U. K.; Stemmler, T. L.; Klishko, V. Y.; Li, S.; Albertine, K. H.; Davis, D. R.; Sundquist, W. I. Proteolytic refolding of the HIV-1 capsid protein amino-terminus facilitates viral core assembly. The EMBO Journal 1998, 17, 1555–1568.

20. Dick, R. A.; Zadrozny, K. K.; Xu, C.; Schur, F. K. M.; Lyddon, T. D.; Ricana, C. L.; Wagner, J. M.; Perilla, J. R.; Ganser-Pornillos, B. K.; Johnson, M. C.; Pornillos, O.; Vogt, V. M. Inositol phosphates are assembly co-factors for HIV-1. Nature 2018, 560, 509–512.

21. Dick, R. A.; Mallery, D. L.; Vogt, V. M.; James, L. C. IP6 Regulation of HIV Capsid Assembly, Stability, and Uncoating. Viruses 2018, 10, 640.

22. Renner, N.; Kleinpeter, A.; Mallery, D. L.; Albecka, A.; Rifat Faysal, K. M.; Böcking, T.; Saiardi, A.; Freed, E. O.; James, L. C. HIV-1 is dependent on its immature lattice to recruit IP6 for mature capsid assembly. Nature Structural & Molecular Biology 2023, 30, 370–382.

23. Mallery, D. L.; Kleinpeter, A. B.; Renner, N.; Faysal, K. M. R.; Novikova, M.; Kiss, L.; Wilson, M. S. C.; Ahsan, B.; Ke, Z.; Briggs, J. A. G.; Saiardi, A.; Böcking, T.; Freed, E. O.; James, L. C. A stable immature lattice packages IP6 for HIV capsid maturation. Science Advances 2021, 7, eabe4716.

24. Rebensburg, S. V.; Wei, G.; Larue, R. C.; Lindenberger, J.; Francis, A. C.; Annamalai, A. S.; Morrison, J.; Shkriabai, N.; Huang, S.-W.; KewalRamani, V.; Poeschla, E. M.; Melikyan, G. B.; Kvaratskhelia, M. Sec24C is an HIV-1 host dependency factor crucial for virus replication. Nature Microbiology 2021, 6, 435–444.

25. Bhattacharya, A.; Alam, S. L.; Fricke, T.; Zadrozny, K.; Sedzicki, J.; Taylor, A. B.; Demeler, B.; Pornillos, O.; Ganser-Pornillos, B. K.; Diaz-Griffero, F.; Ivanov, D. N.; Yeager, M. Structural basis of HIV-1 capsid recognition by PF74 and CPSF6. Proceedings of the National Academy of Sciences 2014, 111, 18625–18630, Publisher: Proceedings of the National Academy of Sciences.

26. Wei, G. et al. Prion-like low complexity regions enable avid virus-host interactions during HIV-1 infection. Nature Communications 2022, 13, 5879.

27. Price, A. J.; Jacques, D. A.; McEwan, W. A.; Fletcher, A. J.; Essig, S.; Chin, J. W.; Halambage, U. D.; Aiken, C.; James, L. C. Host cofactors and pharmacologic ligands share an essential interface in HIV-1 capsid that is lost upon disassembly. PLoS Pathog 2014, 10, e1004459.

28. Blair, W. S. et al. HIV Capsid is a Tractable Target for Small Molecule Therapeutic Intervention. PLOS Pathogens 2010, 6, e1001220, Publisher: Public Library of Science.

29. Link, J. O. et al. Clinical targeting of HIV capsid protein with a long-acting small molecule. Nature 2020, 584, 614–618, Publisher: Nature Publishing Group.

30. Bester, S. M. et al. Structural and mechanistic bases for a potent HIV-1 capsid inhibitor. Science 2020, 370, 360–364, Publisher: American Association for the Advancement of Science.

31. Rankovic, S.; Ramalho, R.; Aiken, C.; Rousso, I. PF74 Reinforces the HIV-1 Capsid To Impair Reverse Transcription-Induced Uncoating. J Virol 2018, 92.

32. Perilla, J. R.; Schulten, K. Physical properties of the HIV-1 capsid from all-atom molecular dynamics simulations. Nature Communications 2017, 8, 15959, Publisher: Nature Publishing Group.

33. Yu, A.; Lee, E. M. Y.; Briggs, J. A. G.; Ganser-Pornillos, B. K.; Pornillos, O.; Voth, G. A. Strain and rupture of HIV-1 capsids during uncoating. Proceedings of the National Academy of Sciences 2022, 119, e2117781119.

34. Craveur, P.; Gres, A. T.; Kirby, K. A.; Liu, D.; Hammond, J. A.; Deng, Y.; Forli, S.; Goodsell, D. S.; Williamson, J. R.; Sarafianos, S. G.; Olson, A. J. Novel Intersubunit Interaction Critical for HIV-1 Core Assembly Defines a Potentially Targetable Inhibitor Binding Pocket. mBio 2019, 10, e02858–18.

35. Gres, A. T.; Kirby, K. A.; McFadden, W. M.; Du, H.; Liu, D.; Xu, C.; Bryer, A. J.; Perilla, J. R.; Shi, J.; Aiken, C.; Fu, X.; Zhang, P.; Francis, A. C.; Melikyan, G. B.; Sarafianos, S. G. Multidisciplinary studies with mutated HIV-1 capsid proteins reveal structural mechanisms of lattice stabilization. Nature Communications 2023, 14, 5614.

36. Yu, N.; Hagan, M. F. Simulations of HIV Capsid Protein Dimerization Reveal the Effect of Chemistry and Topography on the Mechanism of Hydrophobic Protein Association. Biophysical Journal 2012, 103, 1363–1369.

37. Sun, Q.; Levy, R. M.; Kirby, K. A.; Wang, Z.; Sarafianos, S. G.; Deng, N. Molecular Dynamics Free Energy Simulations Reveal the Mechanism for the Antiviral Resistance of the M66I HIV-1 Capsid Mutation. Viruses 2021, 13, 920.

38. Ni, T.; Zhu, Y.; Yang, Z.; Xu, C.; Chaban, Y.; Nesterova, T.; Ning, J.; Böcking, T.; Parker, M. W.; Monnie, C.; Ahn, J.; Perilla, J. R.; Zhang, P. Structure of native HIV-1 cores and their interactions with IP6 and CypA. Science Advances 2021, 7, eabj5715.

39. Yu, A.; Lee, E. M. Y.; Jin, J.; Voth, G. A. Atomic-scale characterization of mature HIV-1 capsid stabilization by inositol hexakisphosphate (IP6). Science Advances 2020, 6, eabc6465.

40. Sha, H.; Zhu, F. Hexagonal Lattices of HIV Capsid Proteins Explored by Simulations Based on a Thermodynamically Consistent Model. The Journal of Physical Chemistry B 2024, pcs.jpcb.3c06881.

41. Gupta, M.; Pak, A. J.; Voth, G. A. Critical mechanistic features of HIV-1 viral capsid assembly. SCIENCE ADVANCES 2023,

42. Grime, J. M. A.; Dama, J. F.; Ganser-Pornillos, B. K.; Woodward, C. L.; Jensen, G. J.; Yeager, M.; Voth, G. A. Coarse-grained simulation reveals key features of HIV-1 capsid self-assembly. Nature Communications 2016, 7, 11568.

43. Qiao, X.; Jeon, J.; Weber, J.; Zhu, F.; Chen, B. Mechanism of polymorphism and curvature of HIV capsid assemblies probed by 3D simulations with a novel coarse grain model. Biochimica et Biophysica Acta (BBA) - General Subjects 2015, 1850, 2353–2367.

44. Qiao, X.; Jeon, J.; Weber, J.; Zhu, F.; Chen, B. Construction of a novel coarse grain model for simulations of HIV capsid assembly to capture the backbone structure and inter-domain motions in solution. Data in Brief 2015, 5, 506–512.

45. Chen, B.; Tycko, R. Simulated Self-Assembly of the HIV-1 Capsid: Protein Shape and Native Contacts Are Sufficient for Two-Dimensional Lattice Formation. Biophysical Journal 2011, 100, 3035–3044.

46. Krishna, V.; Ayton, G. S.; Voth, G. A. Role of protein interactions in defining HIV-1 viral capsid shape and stability: a coarse-grained analysis. Biophys J 2010, 98, 18–26.

47. Yu, Z.; Dobro, M. J.; Woodward, C. L.; Levandovsky, A.; Danielson, C. M.; Sandrin, V.; Shi, J.; Aiken, C.; Zandi, R.; Hope, T. J.; Jensen, G. J. Unclosed HIV-1 Capsids Suggest a Curled Sheet Model of Assembly. Journal of Molecular Biology 2013, 425, 112–123.

48. Jennings, J.; Shi, J.; Varadarajan, J.; Jamieson, P. J.; Aiken, C. The Host Cell Metabolite Inositol Hexakisphosphate Promotes Efficient Endogenous HIV-1 Reverse Transcription by Stabilizing the Viral Capsid. mBio 2020, 11.

49. Mallery, D. L.; Faysal, K. M. R.; Kleinpeter, A.; Wilson, M. S. C.; Vaysburd, M.; Fletcher, A. J.; Novikova, M.; Bocking, T.; Freed, E. O.; Saiardi, A.; James, L. C. Cellular IP6 Levels Limit HIV Production while Viruses that Cannot Efficiently Package IP6 Are Attenuated for Infection and Replication. Cell Rep 2019, 29, 3983–3996 e4.

50. Rankovic, S.; Varadarajan, J.; Ramalho, R.; Aiken, C.; Rousso, I. Reverse Transcription Mechanically Initiates HIV-1 Capsid Disassembly. J Virol 2017, 91.

51. Marquez, C. L.; Lau, D.; Walsh, J.; Shah, V.; McGuinness, C.; Wong, A.; Aggarwal, A.; Parker, M. W.; Jacques, D. A.; Turville, S.; Bocking, T. Kinetics of HIV-1 capsid uncoating revealed by single-molecule analysis. Elife 2018, 7 .

52. Schirra, R. T.; dos Santos, N. F. B.; Zadrozny, K. K.; Kucharska, I.; Ganser-Pornillos, B. K.; Pornillos, O. A molecular switch modulates assembly and host factor binding of the HIV-1 capsid. Nature Structural & Molecular Biology 2023, 30, 383–390.

53. Kobayakawa, T.; Yokoyama, M.; Tsuji, K.; Fujino, M.; Kurakami, M.; Boku, S.; Nakayama, M.; Kaneko, M.; Ohashi, N.; Kotani, O.; Murakami, T.; Sato, H.; Tamamura, H. Small-Molecule Anti-HIV-1 Agents Based on HIV-1 Capsid Proteins. Biomolecules 2021, 11, 208.

54. Humphrey, W.; Dalke, A.; Schulten, K. VMD: Visual molecular dynamics. Journal of Molecular Graphics 1996, 14, 33–38.

55. Highland, C. M.; Tan, A.; Ricaña, C. L.; Briggs, J. A. G.; Dick, R. A. Structural insights into HIV-1 polyanion-dependent capsid lattice formation revealed by single particle cryo-EM. Proceedings of the National Academy of Sciences 2023, 120, e2220545120, Publisher: Proceedings of the National Academy of Sciences.

56. Dolinsky, T. J.; Nielsen, J. E.; McCammon, J. A.; Baker, N. A. PDB2PQR: an automated pipeline for the setup of Poisson-Boltzmann electrostatics calculations. Nucleic Acids Res 2004, 32, W665–7.

57. Huang, J.; Rauscher, S.; Nawrocki, G.; Ran, T.; Feig, M.; de Groot, B. L.; Grubmuller, H.; MacKerell, A. D. CHARMM36m: an improved force field for folded and intrinsically disordered proteins. Nat Methods 2017, 14, 71–73.

58. Beglov, D.; Roux, B. Finite representation of an infinite bulk system: Solvent boundary potential for computer simulations. The Journal of Chemical Physics 1994, 100, 9050–9063.

59. Jorgensen, W. L.; Chandrasekhar, J.; Madura, J. D.; Impey, R. W.; Klein, M. L. Comparison of simple potential functions for simulating liquid water. Journal of Chemical Physics 1983, 79, 926.

60. Vanommeslaeghe, K.; Hatcher, E.; Acharya, C.; Kundu, S.; Zhong, S.; Shim, J.; Darian, E.; Guvench, O.; Lopes, P.; Vorobyov, I.; Mackerell, A. D. CHARMM general force field: A force field for drug-like molecules compatible with the CHARMM all-atom additive biological force fields. Journal of Computational Chemistry 2010, 31, 671–690.

61. Phillips, J. C.; Braun, R.; Wang, W.; Gumbart, J.; Tajkhorshid, E.; Villa, E.; Chipot, C.; Skeel, R. D.; Kale, L.; Schulten, K. Scalable molecular dynamics with NAMD. J Comput Chem 2005, 26, 1781–802.

62. Phillips, J. C. et al. Scalable molecular dynamics on CPU and GPU architectures with NAMD. The Journal of Chemical Physics 2020, 153, 044130.

63. Adelman, S. A.; Doll, J. D. Generalized langevin equation approach for atom/solid-surface scattering: collinear atom/harmonic chain model. The Journal of Chemical Physics 1974, 61, 4242–4245.

64. Feller, S. E.; Zhang, Y.; Pastor, R. W.; Brooks, B. R. Constant pressure molecular dynamics simulation: The Langevin piston method. The Journal of Chemical Physics 1995, 103, 4613–4621.

65. Martyna, G. J.; Tobias, D. J.; Klein, M. L. Constant pressure molecular dynamics algorithms. The Journal of Chemical Physics 1994, 101, 4177–4189.

66. Darden, T.; York, D.; Pedersen, L. Particle mesh Ewald: An Nlog(N) method for Ewald sums in large systems. The Journal of Chemical Physics 1993, 98, 10089–10092.

67. Grossfield, A. WHAM: an implementation of the weighted histogram analysis method, version 2.0.10.

68. Bouysset, C.; Fiorucci, S. ProLIF: a library to encode molecular interactions as fingerprints. Journal of Cheminformatics 2021, 13, 72.

69. Michaud-Agrawal, N.; Denning, E. J.; Woolf, T. B.; Beckstein, O. MDAnalysis: A toolkit for the analysis of molecular dynamics simulations. Journal of Computational Chemistry 2011, 32, 2319–2327.

