## Supplementary Information for "IP6 and PF74 affect HIV-1 Capsid Stability through Modulation of Hexamer-Hexamer Tilt Angle Preference"

#### Additional System Preparation Notes

##### Choosing a Hexamer-Pentamer Pair

The two whole-capsid models (3J3Q and 3J3Y)[(1)] contain 12 pentamers which each form 5 interfaces with surrounding hexamers for a grand total of 120 HP interfaces to choose from. In order to narrow down this list, we first calculated each HP pair's tilt and twist angles using the average plane produced by the centers of mass of each of the 5 or 6 chains making up a pentamer or hexamer, respectively. Previous studies that reported the normal ranges of tilt and twist angles were not broken down into HH and HP groups, however, it was to be expected that the HP interfaces would have higher tilt angles due to the pentamer's dome-like structure as well as the theoretical purpose of pentamers in the fullerene cone complex to induce local curvature and assist in closing of the ends. Thus, unsurprisingly, we found that the range of tilt angles found in the HP interfaces was narrower and higher Figure 1.

In addition to tilt/twist angles, the various HP pairs in the two whole-capsid models also vary in the non-bonded interactions present at the inter-oligomer interface. Since the available softwares require hydrogens to be present in order to evaluate hydrogen bonding, we built a fully protonated version of each whole capsid and minimized the structures for 10,000 steps via steepest descent to reduce any clashes produced by poorly-guessed hydrogen positions. From this minimized structure, we used the PROLIF package to assess each interface for non-bonded interactions (hydrogen bonds, salt bridges, and pi-pi stacks), and then generated a list of all the interactions that were found. Then for each interface, we normalized the number of intact interactions against the total to come up with a score for each HP pair. We then generated a list of the interfaces that had tilt/twist values close to the average (for HP interfaces), which we coarsely set to a range of 25-32° tilt and <5° twist, AND had a composite score close to the average.

The chosen HP pair had tilt/twist values of 27.5°/1.8°. It is composed of the following segment ID's from the 3J3Y structure: (UC, FD, ME, XE, QD, BE) and (TXA, UXA, VXA, WXA, XXA)

#### Preparation of the IP6-containing Hexamer-Hexamer System

The 6BHS[(2)] structure was used with the following mutations: Mutations made due to variations in sequence between 3J34 and 6BHS:
A92E
G208A
Experimental mutations in 6BHS:
A184W
A185M
C45E
C14A
Missing residues in 6BHS:
88-89
177-186
219-end

#### Histidine Protonations

Histidine protonations are in Table 1 and were determined using the PROPKA[(3)] online server with modifications in the following instances for consistency between protein chains:
**His12** on chain F was made to be HSD but on inspection, this His is rotated upward in the model whereas His12 is rotated downward in all other chains, making a favorable interaction with Asp51. His12 on chain F was not making favorable interactions with anything so we switched this residue to HSE so that it will have the opportunity to interact with Asp51 in simulation.
**His87** on chains A and J were modeled as HSP due to formation of a salt bridge with Glu98. His87 on chains F and I were modeled as HSD because they're rotated away from Glu98 but we switched them to HSP to allow it to rotate and form favorable interactions with Glu98 in simulation.
**His62** is at the interface between the two subunits from the same hexamer and in two chains was pointing at the backbone oxygen of Glu28. The other two were modeled as HSD and were not engaging in favorable hydrogen bonding so we switched them to HSE.

#### Equilibration Protocol

Varying parameters for each step of the equilibration are listed in Table 2. All other parameters are the same between steps.

#### SMD Restraints

**Breaking Interface:** Five backbone carbons from the chains A and F were restrained by a harmonic restraint of 1 kcal/mol/Å$^2$ in the X dimension to prevent the hexamer centered at the origin from drifting during pulling (Table 3)

**Table 1.** Histidine protonation states common to all simulations. HSE designates histidine with the epsilon nitrogen protonated, HSD designates histidine with the delta nitrogen protonated, and HSP designates histidine with both delta and epsilon nitrogens protonated. Histidine protonation states were predicted with PROPKA and selectively altered to reflect consistency between protein chains.

| Residue | Histidine Protonation |
|---|---|
| 12 | HSE |
| 62 | HSE |
| 84 | HSD |
| 87 | HSP |
| 120 | HSD |
| 226 | HSD |

**Table 2.** Simulated annealing protocol for system equilibration.

| Stage | Steps | Timestep (fs/s) | Time | Protein Backbone Restraint (kcal/mol/Å$^2$) | Pressure | Reassign Temperature |
|---|---|---|---|---|---|---|
| 1 | 25,000 | 1 | 25 ps | 10 | OFF | ON |
| 2 | 25,000 | 1 | 25 ps | 5 | OFF | ON |
| 3 | 25,000 | 1 | 25 ps | 2.5 | ON | ON |
| 4 | 100,000 | 2 | 200 ps | 1 | ON | ON |
| 5 | 100,000 | 2 | 200 ps | 0.5 | ON | ON |
| 6 | 49,525,000 | 2 | 99.05 ns | 0.5 | ON | OFF |

**Table 3.** Backbone atoms restrained in cartesian space during interface breaking SMD simulations to prevent drift of the hexamer starting at the origin.

| Residue Name | Residue Number | Atom Type | Chain |
|---|---|---|---|
| LEU | 202 | CA | A |
| THR | 216 | CA | A |
| ARG | 154 | CA | F |
| THR | 186 | CA | F |
| ILE | 201 | CA | F |

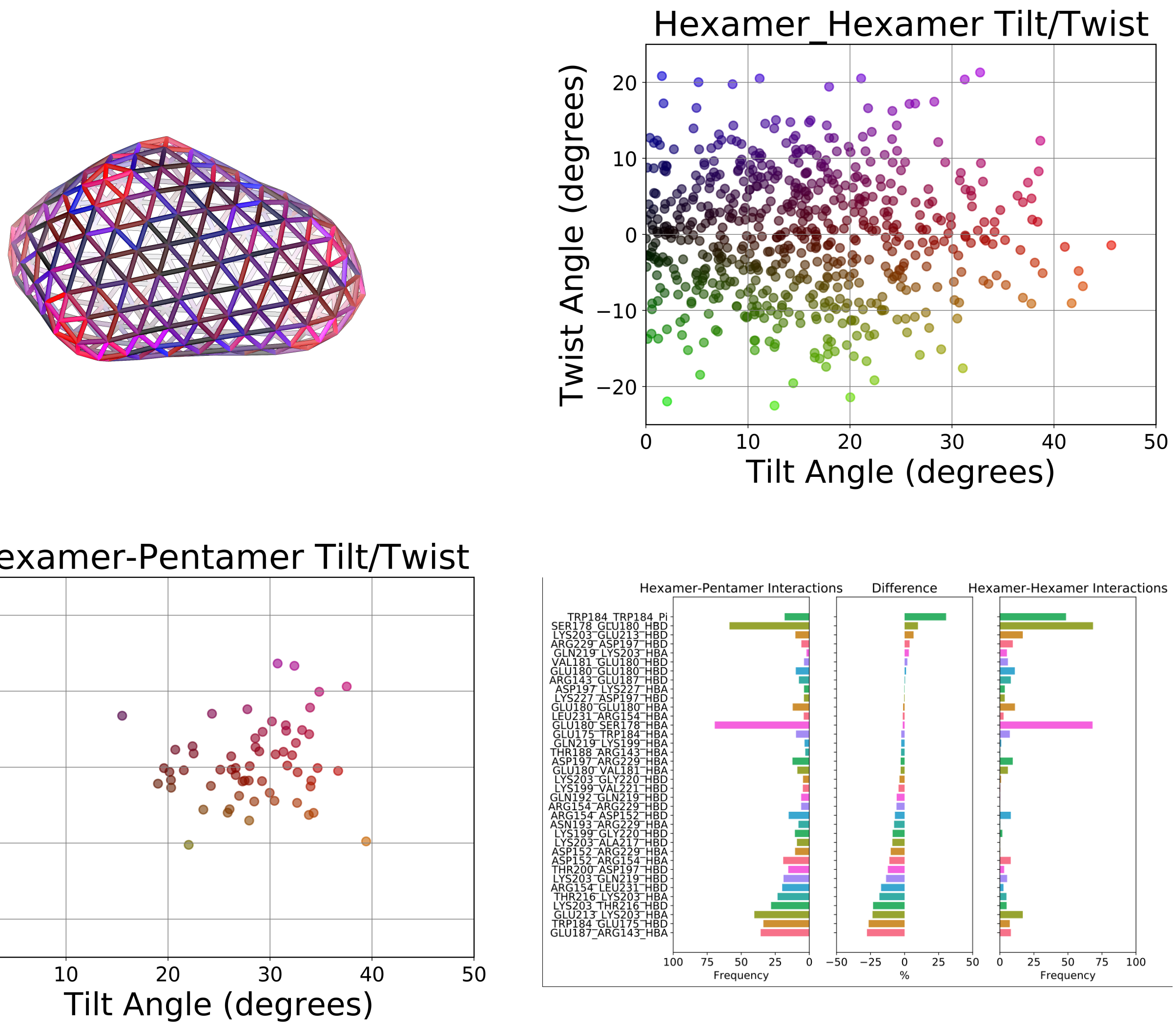


**Fig. 1.** Top left: wirecage model of a whole capsid with sticks drawn from the center of mass of a hexamer or pentamer group to each of its neighbors. Sticks are color coded by curvature with red saturation representing the tilt angle and blue(+)/green(-) saturation representing the twist angle between two neighboring groups. Thus, black is low-tilt and low-twist, red is high-tilt and low-twist, blue is low-tilt and high-positive-twist, purple is high-tilt and high-postive-twist, green is low-tilt and high-negative-twist, and yellow is high-tilt and high-negative-twist. Top right is the distribution of curvatures found between all hexamer-hexamer pairs and bottom left is the distribution of curvatures between all hexamer-pentamer pairs. Scatter plots share the same coloring scheme as the wirecage model. In the bottom left is an analysis of the non-bonded interactions found at the various interfaces between neighboring groups, sectioned into hexamer-hexamer and hexamer-pentamer interfaces on the sides with the difference between the two in the middle. All non-bonded interactions data and tilt/twist angles are collated between the PDB:3J3Q and 3J3Y structures.

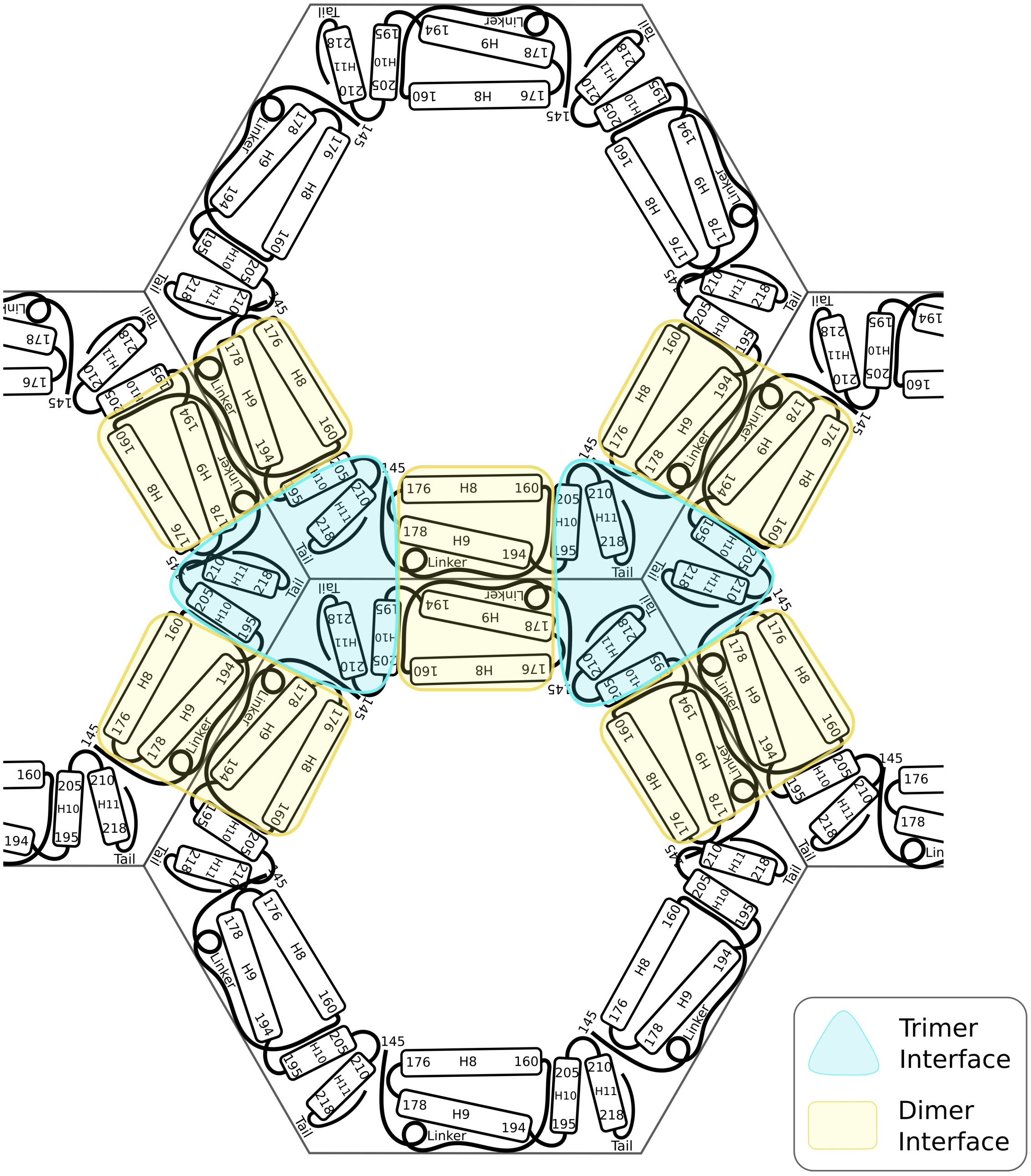


**Fig. 2.** Model of the secondary structural features involved in the dimer and trimer interfaces. Only the C-terminal domain is shown for each CA monomer for simplicity, where the N-terminal domains would connect at residue 145. Helices are labeled with "H" and the first and last residue in a helix are shown by residue number.

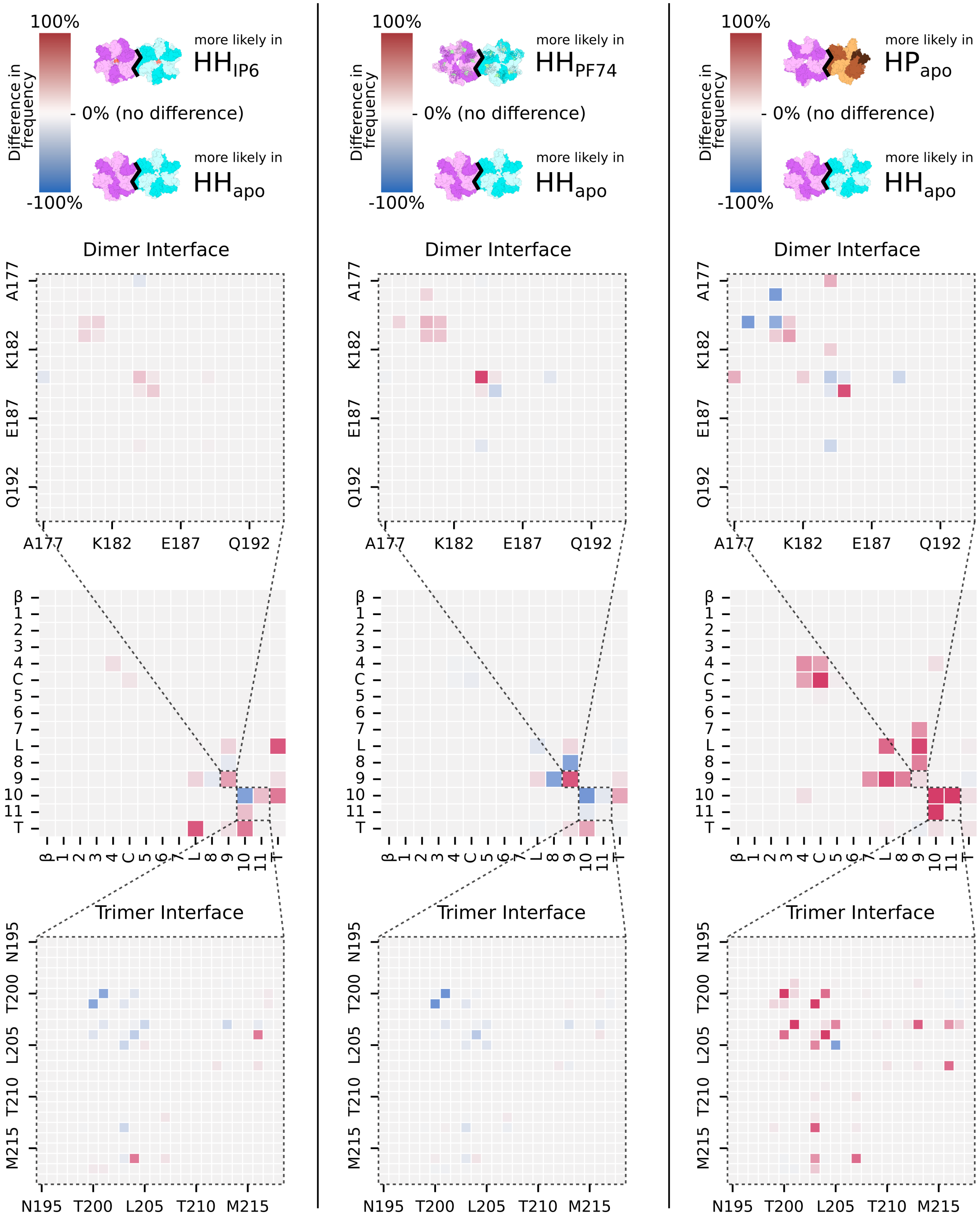


**Fig. 3.** Difference in non-bonded interaction frequencies at the inter-oligomer interface for $HH_{IP6}$ (left), $HH_{PF74}$ (middle), and $HP_{apo}$ (right), with respect to $HH_{apo}$, averaged across all simulations. First row: Keys for each column. Second row: differences in interactions between helix 9 (representative of the dimer interface) from each of the opposing oligomers, in order of the amino acid sequence. Third row: Collective difference in frequencies by secondary structural features as defined by Figure 1D. Fourth row: heatmap of differences at the trimer interface, including helices 10 and 11.

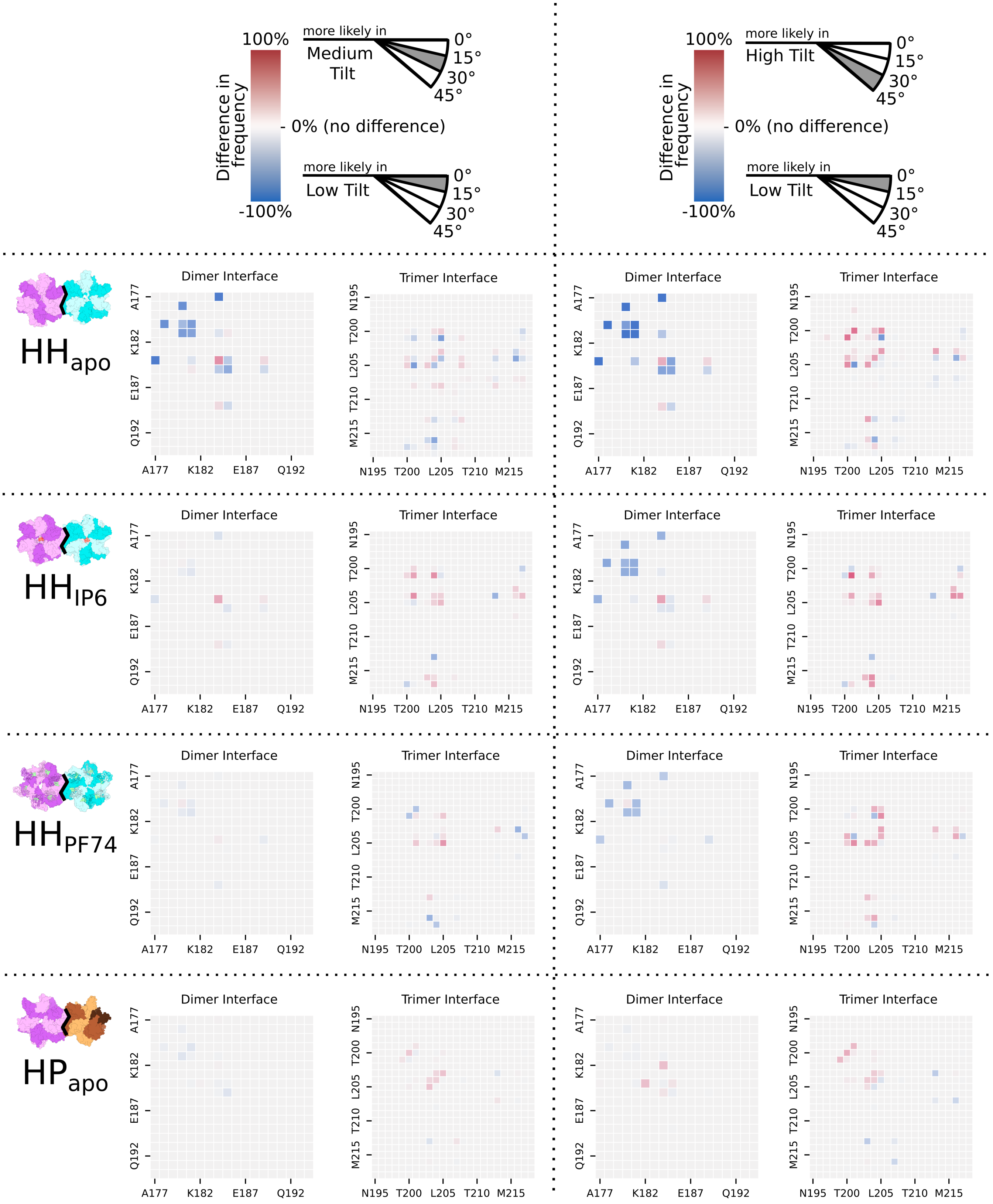


**Fig. 4.** Difference in non-bonded interaction frequency at the HH or HP inter-oligomer interface for $HH_{apo}$, $HH_{IP6}$, $HH_{PF74}$, and $HP_{apo}$ systems at middle- (left panel) and high-tilt (right panel) angles, with respect to themselves at low tilt angles. Residues in sequential order are plotted from helix 9, representative of the dimer interface, and helices 10 and 11, representative of the trimer interface.

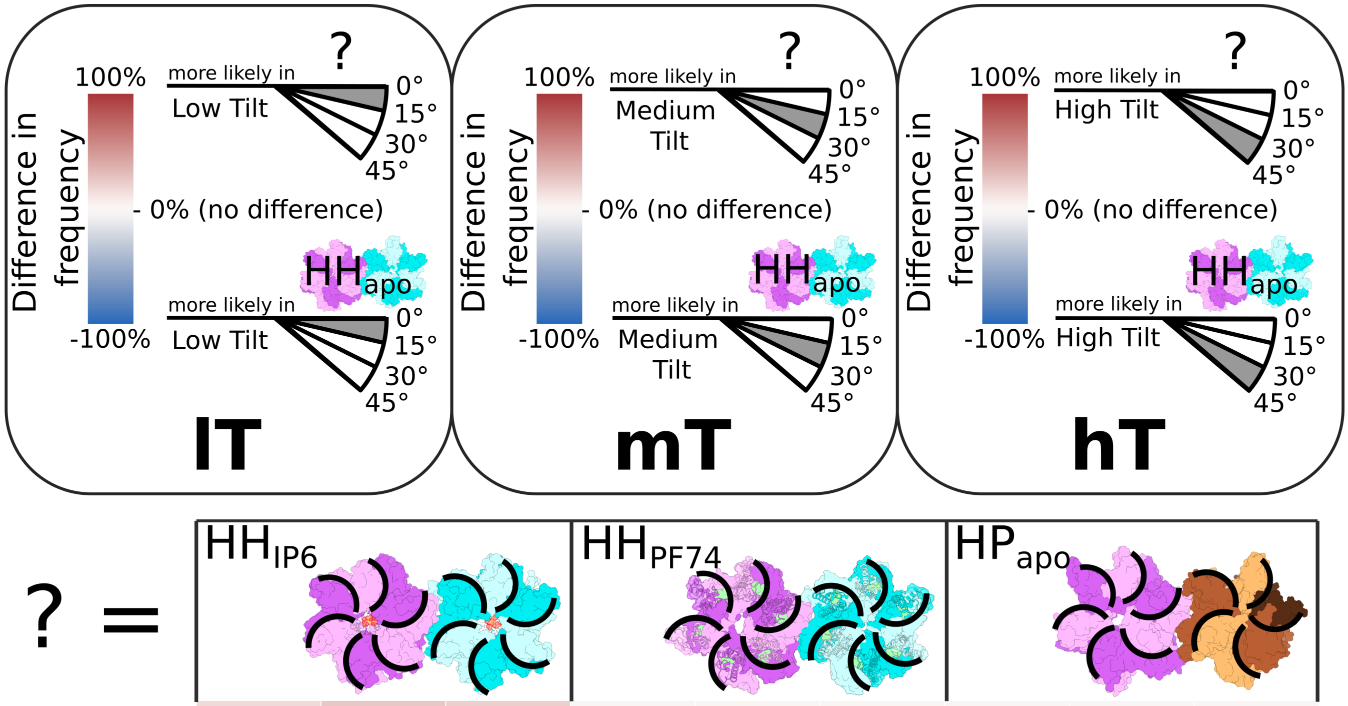


| | IP6 lT | IP6 mT | IP6 hT | PF74 lT | PF74 mT | PF74 hT | HP lT | HP mT | HP hT |
|---|---|---|---|---|---|---|---|---|---|
| N5 -H- L6 | 16.7 | 25.8 | 22.0 | -2.1 | 1.7 | 1.6 | -1.4 | -1.0 | -0.8 |
| N5 )( L6 | 18.0 | 25.5 | 24.9 | -3.7 | 1.4 | 2.2 | -1.3 | -0.3 | 1.0 |
| L6 )( L6 | 26.8 | 4.5 | 5.8 | 0.2 | -1.7 | -2.2 | -1.2 | -3.1 | -2.6 |
| P38 )( N57 | 6.9 | 7.6 | 7.3 | 10.0 | 13.5 | 12.7 | -22.2 | -20.1 | -20.4 |
| N57 -H- R173 | 24.7 | 29.3 | 26.4 | 27.5 | 36.1 | 37.0 | -7.2 | -7.4 | -8.6 |
| V59 -H- R173 | 25.0 | 27.9 | 25.2 | 26.0 | 34.4 | 34.7 | -9.7 | -6.7 | -6.8 |
| A64 )( Y169 | | | -11.3 | | | -21.3 | | | 1.6 |

**Fig. 5.** Difference in frequency of non-bonded interactions present at inter-oligomer interface of $HH_{IP6}$, $HH_{PF74}$, and $HP_{apo}$, with respect to $HH_{apo}$, at low, medium, or high-tilt ranges. Values fall into the range from +100% to -100% where zero indicates there is no difference between the systems, but not necessarily that the interaction was absent.

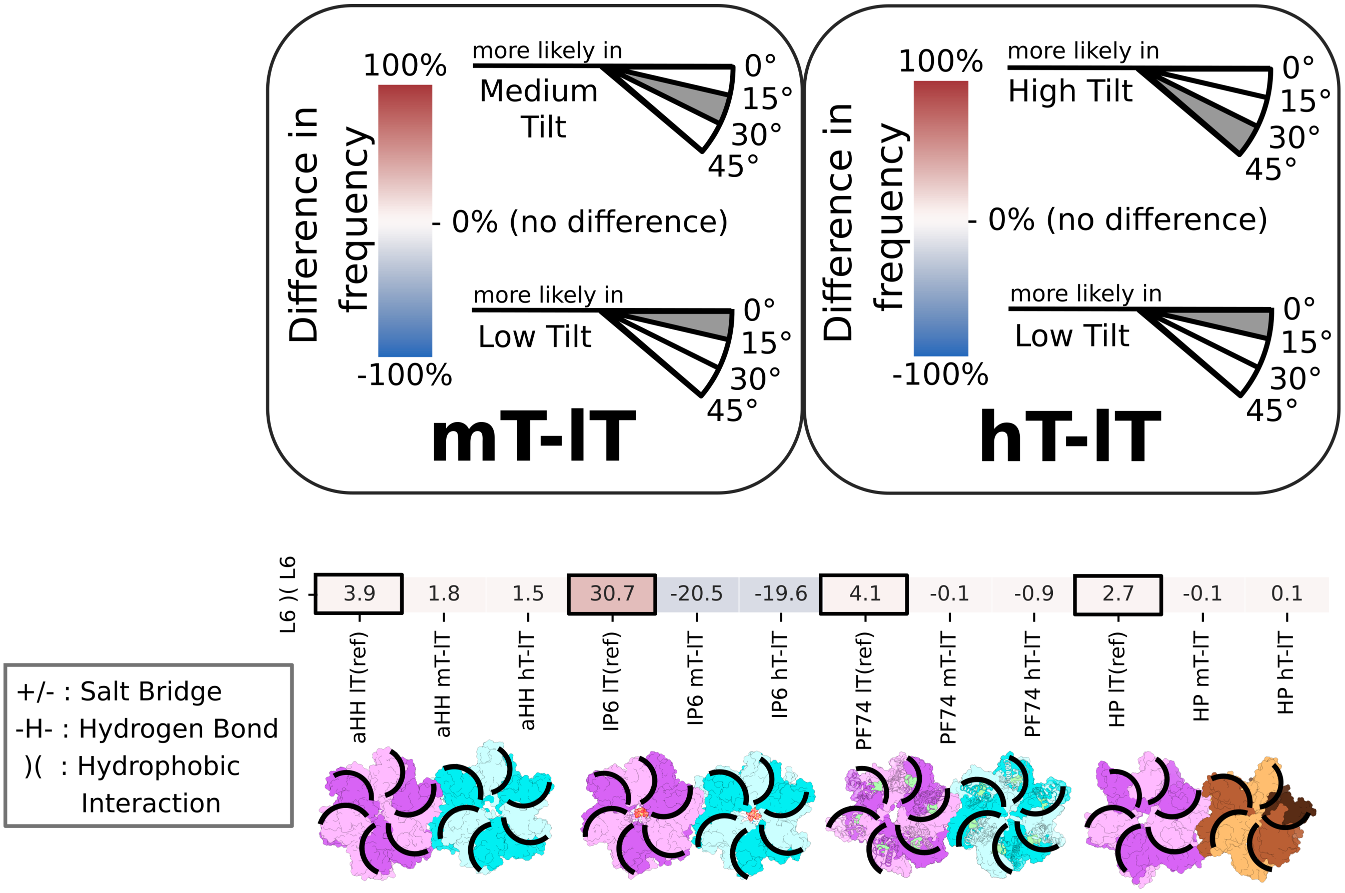


**Fig. 6.** Difference in frequency of non-bonded interactions present between capsid monomers (intra-oligomer interface) in the $HH_{IP6}$, $HH_{PF74}$, and $HP_{apo}$ systems, with respect to intra-oligomer interfaces in the $HH_{apo}$ system, at low, medium, or high-tilt ranges.

| | aHH lT(ref) | aHH mT-lT | aHH hT-lT | IP6 lT(ref) | IP6 mT-lT | IP6 hT-lT | PF74 lT(ref) | PF74 mT-lT | PF74 hT-lT | HP lT(ref) | HP mT-lT | HP hT-lT |
|---|---|---|---|---|---|---|---|---|---|---|---|---|
| E79 +/– R82 | 0.0 | -0.0 | -0.0 | 0.0 | 0.0 | 0.0 | 0.1 | -0.1 | -0.1 | 32.0 | -28.8 | -31.9 |
| L83 -H- V86 | 0.0 | 0.0 | 0.0 | 0.0 | 0.0 | 0.0 | 0.0 | 0.0 | 0.0 | 31.0 | -31.0 | -31.0 |
| L83 )( V86 | 0.7 | -0.7 | -0.7 | 0.0 | 0.0 | 0.0 | 0.0 | 0.0 | 0.0 | 45.9 | -43.1 | -45.9 |
| H84 )( V86 | 0.0 | 0.0 | 0.0 | 0.0 | 0.0 | 0.0 | 0.0 | 0.0 | 0.0 | 22.6 | -22.5 | -22.6 |
| P85 )( P85 | 0.8 | -0.8 | -0.8 | 0.0 | 0.0 | 0.0 | 0.0 | 0.0 | 0.0 | 35.3 | -34.7 | -35.3 |
| P147 -H- Q192 | 3.8 | 6.0 | 28.5 | 0.5 | 0.1 | -0.4 | 0.0 | 0.0 | 0.2 | 0.0 | 0.0 | 0.0 |
| I150 )( W184 | 62.3 | -20.9 | -17.2 | 54.8 | -11.7 | -36.5 | 31.5 | -10.3 | -26.4 | 73.1 | -5.3 | -0.3 |
| I150 )( L189 | 22.7 | -19.3 | -22.7 | 3.0 | -2.9 | -3.0 | 0.0 | 0.0 | 0.0 | 1.1 | 0.7 | 3.2 |
| L151 )( L151 | 55.4 | -2.2 | 12.4 | 45.0 | 4.2 | 10.1 | 38.3 | 11.3 | 21.9 | 7.9 | 9.0 | 8.2 |
| L151 )( W184 | 67.2 | -8.3 | -21.5 | 64.5 | 7.3 | -1.6 | 73.2 | -1.0 | -3.8 | 1.9 | 1.8 | 0.6 |
| L151 )( T188 | 14.2 | 2.9 | -3.5 | 15.7 | 13.9 | 9.0 | 36.5 | -4.2 | -28.2 | 39.3 | -1.5 | -1.7 |
| L151 )( Q192 | 32.4 | 25.6 | 4.1 | 57.1 | 4.5 | 1.2 | 65.7 | -9.6 | -37.1 | 73.5 | -0.7 | -2.0 |
| D152 +/– R154 | 0.0 | 22.8 | 52.3 | 0.2 | 0.8 | 10.0 | 0.9 | 0.7 | 12.9 | 59.6 | 4.0 | 7.8 |
| D152 -H- Q192 | 1.2 | 18.9 | 55.7 | 0.3 | 1.0 | 0.0 | 2.7 | 0.3 | -2.3 | 0.0 | 0.0 | 0.2 |
| Q155 -H- R229 | 0.0 | 0.0 | 0.0 | 0.0 | 11.6 | 26.8 | 0.0 | 0.0 | 0.0 | 0.0 | 0.0 | 0.0 |
| P157 )( R229 | 0.0 | 0.0 | 0.0 | 0.1 | 17.4 | 36.9 | 0.0 | 0.0 | 0.0 | 0.0 | 0.0 | 0.0 |
| L172 )( W184 | 42.3 | -39.5 | -42.3 | 12.7 | -12.4 | -12.7 | 0.0 | 0.0 | 0.0 | 38.8 | -11.6 | -8.3 |
| E175 )( W184 | 64.0 | -53.2 | -62.4 | 41.7 | -9.7 | -39.0 | 1.0 | -0.3 | -1.0 | 73.1 | 0.7 | -1.9 |
| A177 )( W184 | 80.5 | -46.9 | -73.2 | 41.8 | -7.0 | -25.6 | 51.6 | -0.6 | -15.8 | 74.2 | 0.6 | 0.5 |
| S178 -H- E180 | 63.4 | -39.4 | -60.8 | 43.8 | 0.5 | -29.9 | 55.5 | -2.8 | -22.5 | 5.8 | -2.2 | -1.9 |
| E180 )( E180 | 37.5 | -22.4 | -33.9 | 46.8 | 0.8 | -24.4 | 43.8 | 2.8 | 3.2 | 0.1 | -0.0 | 0.1 |
| E180 )( V181 | 63.4 | -34.3 | -58.2 | 58.3 | -2.2 | -27.7 | 62.6 | -3.3 | -21.2 | 61.9 | -5.4 | -0.8 |
| V181 )( V181 | 69.4 | -31.8 | -63.2 | 53.2 | -5.0 | -28.1 | 66.7 | -5.3 | -20.1 | 72.6 | -0.9 | -0.8 |
| W184 )( W184 | 7.1 | 27.1 | 18.8 | 43.9 | 20.2 | 21.6 | 66.6 | 2.6 | 0.6 | 4.1 | -1.6 | -1.3 |
| W184 )( M185 | 82.7 | -16.1 | -31.4 | 74.8 | 0.3 | -5.4 | 74.6 | 0.2 | -0.8 | 62.0 | -0.4 | 4.1 |
| M185 )( M185 | 30.8 | -28.1 | -30.8 | 5.9 | -5.8 | -5.9 | 0.2 | -0.2 | -0.2 | 60.1 | -5.5 | -2.3 |

+/- : Salt Bridge
-H- : Hydrogen Bond
)( : Hydrophobic Interaction

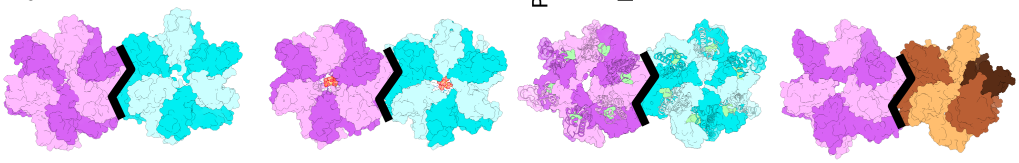

**Fig. 7.** Difference in frequency of non-bonded interactions present at the inter-oligomer interface in the $HH_{IP6}$, $HH_{PF74}$, and $HP_{apo}$ systems, with respect to inter-oligomer interface in the $HH_{apo}$ system, at low, medium, or high-tilt ranges.

| | aHH lT(ref) | aHH mT-lT | aHH hT-lT | IP6 lT(ref) | IP6 mT-lT | IP6 hT-lT | PF74 lT(ref) | PF74 mT-lT | PF74 hT-lT | HP lT(ref) | HP mT-lT | HP hT-lT |
|---|---|---|---|---|---|---|---|---|---|---|---|---|
| Q192 )( L231 | 21.1 | -20.3 | -20.5 | 0.5 | -0.5 | -0.5 | 2.2 | -0.0 | 8.1 | 0.0 | 0.0 | 0.0 |
| N195 -H- R229 | 0.0 | 0.0 | 0.1 | 0.0 | 16.8 | 31.9 | 0.0 | 0.0 | 0.0 | 0.0 | 0.0 | 0.0 |
| P196 )( Q219 | 0.0 | 4.8 | 36.9 | 0.0 | 0.0 | 0.0 | 0.0 | 0.0 | 0.0 | 0.0 | 0.1 | 0.0 |
| P196 )( H226 | 0.0 | 0.3 | 1.2 | 0.9 | 23.3 | 25.2 | 0.0 | 0.0 | 0.0 | 0.0 | 0.0 | 3.0 |
| P196 )( V230 | 0.0 | 7.1 | 34.4 | 0.2 | 1.8 | -0.1 | 0.0 | 0.0 | 0.0 | 0.0 | 0.0 | 0.0 |
| K199 +/– L231 | 0.8 | 33.0 | 51.6 | 4.7 | -4.7 | -4.7 | 24.2 | 5.5 | 25.0 | 0.0 | 0.0 | 0.0 |
| T200 )( I201 | 53.3 | -2.5 | 34.6 | 40.1 | 7.3 | -9.8 | 34.2 | -18.4 | -2.7 | 65.9 | -2.3 | 2.5 |
| T200 )( L205 | 0.3 | 9.7 | 27.9 | 1.7 | -0.8 | 1.7 | 0.0 | 0.1 | 10.3 | 2.0 | 1.8 | 0.5 |
| T200 )( Q219 | 2.4 | 0.3 | 0.6 | 0.2 | 5.9 | 29.5 | 0.3 | -0.3 | 4.3 | 0.0 | 0.0 | 0.0 |
| T200 )( V221 | 9.5 | 18.9 | 31.3 | 2.2 | 5.2 | 11.9 | 0.0 | 0.2 | 1.0 | 0.0 | 0.0 | 1.3 |
| I201 )( I201 | 60.0 | -13.8 | 2.2 | 29.6 | 21.3 | 40.1 | 36.6 | 4.5 | 2.8 | 0.0 | 0.0 | 0.0 |
| I201 )( A204 | 83.9 | -6.2 | 4.4 | 43.9 | 26.7 | 19.9 | 63.5 | 2.3 | -20.9 | 55.4 | 1.1 | 2.3 |
| I201 )( L205 | 60.4 | -33.9 | -36.1 | 47.9 | -5.6 | 2.6 | 24.7 | 11.7 | 27.5 | 0.0 | 0.0 | 0.0 |
| K203 )( L205 | 5.8 | 11.1 | 23.8 | 0.0 | 1.5 | 1.5 | 0.0 | 1.6 | 21.0 | 47.7 | 12.9 | 5.6 |
| K203 +/– E213 | 9.5 | 3.5 | 21.7 | 0.0 | 0.2 | 0.0 | 0.1 | 8.9 | 14.0 | 70.1 | -6.5 | -15.3 |
| K203 )( T216 | 57.5 | -11.1 | 8.3 | 54.4 | 10.1 | 15.5 | 29.3 | -25.7 | 8.6 | 0.5 | 0.9 | 1.5 |
| K203 +/– L231 | 31.6 | -0.3 | -27.2 | 6.7 | -6.7 | -6.7 | 38.3 | 23.1 | 10.4 | 0.0 | 0.0 | 0.0 |
| A204 )( A204 | 18.4 | 9.7 | 22.8 | 1.7 | 8.6 | 4.5 | 5.3 | -4.2 | 1.8 | 98.3 | 9.0 | 9.7 |
| A204 )( E213 | 11.0 | -10.1 | -9.9 | 30.2 | -25.1 | -19.3 | 1.3 | -0.7 | 3.1 | 0.0 | 0.0 | 0.0 |
| A204 )( T216 | 40.8 | -30.0 | -27.1 | 29.8 | 6.2 | 27.0 | 3.5 | -1.5 | 19.5 | 0.0 | 0.0 | 0.0 |
| A204 )( A217 | 62.8 | -7.7 | 8.2 | 39.6 | 14.7 | 25.0 | 47.7 | -17.2 | -12.7 | 0.3 | -0.2 | -0.3 |
| L205 )( L205 | 41.1 | 0.5 | -2.2 | 9.9 | 15.6 | 29.1 | 40.3 | 25.3 | -2.4 | 0.0 | 0.0 | 0.0 |

+/- : Salt Bridge
-H- : Hydrogen Bond
)( : Hydrophobic Interaction

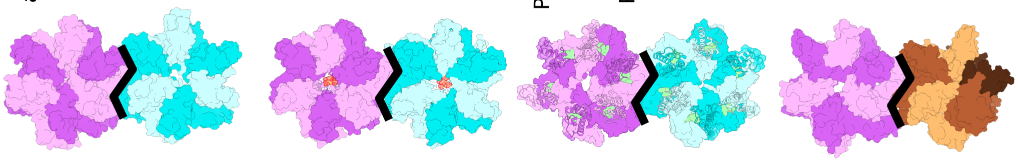

**Fig. 8.** Difference in frequency of non-bonded interactions present at the inter-oligomer interface in the $HH_{IP6}$, $HH_{PF74}$, and $HP_{apo}$ systems, with respect to inter-oligomer interface in the $HH_{apo}$ system, at low, medium, or high-tilt ranges (continued from previous page).

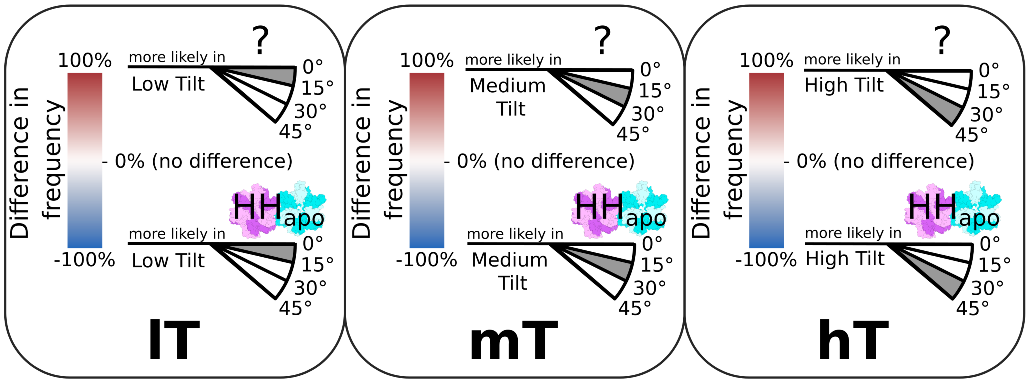


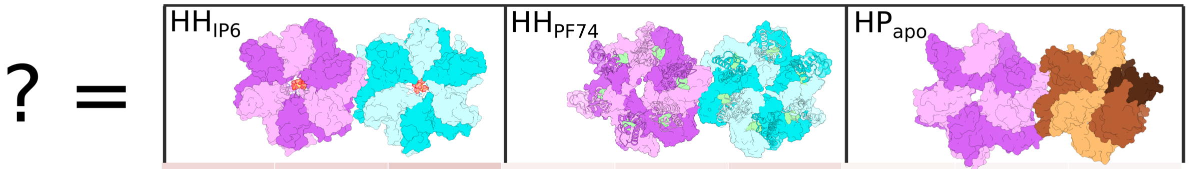


| | IP6 lT | IP6 mT | IP6 hT | PF74 lT | PF74 mT | PF74 hT | HP lT | HP mT | HP hT |
|---|---|---|---|---|---|---|---|---|---|
| I2 )( T119 | 17.5 | 18.7 | 23.5 | 5.5 | 9.0 | 13.6 | 1.9 | -0.4 | -0.2 |
| A22 )( L43 | 29.8 | 27.5 | 27.4 | 18.1 | 12.9 | 12.7 | 31.8 | 25.3 | 26.0 |
| F32 )( Y145 | -12.4 | -10.4 | -14.4 | -17.7 | -21.6 | -21.8 | 9.7 | 6.4 | 6.4 |
| V36 )( L138 | -8.6 | -7.7 | -13.1 | -19.1 | -24.6 | -26.7 | 14.2 | 10.3 | 5.5 |
| F40 )( K131 | -5.4 | -3.0 | | -21.2 | -17.7 | | 1.1 | 1.2 | |
| L56 )( I73 | | -3.9 | -2.8 | | -28.2 | -27.3 | | 1.7 | 1.0 |
| M68 )( Y145 | 13.5 | 11.5 | 12.4 | 21.5 | 26.0 | 27.7 | 0.1 | 0.9 | 2.8 |
| R100 +/– D103 | -4.4 | -1.4 | -4.1 | 18.8 | 22.1 | 20.1 | 10.8 | 11.8 | 10.2 |
| Q155 )( N195 | 0.0 | -4.7 | | 5.0 | 1.1 | | -24.8 | -25.8 | |
| F161 )( M215 | 37.0 | 39.1 | 40.6 | 43.8 | 43.9 | 44.5 | 11.4 | 13.4 | 14.3 |
| D163 -H- R229 | -4.5 | 1.2 | 2.3 | 18.4 | 26.5 | 27.9 | 3.7 | 11.1 | 11.3 |
| V165 )( L211 | 14.1 | 15.3 | 14.6 | 19.8 | 24.7 | 25.5 | 9.3 | 11.2 | 10.8 |
| Y169 -H- T186 | 34.4 | 37.9 | 36.6 | 19.5 | 24.4 | 21.9 | 1.0 | 2.4 | 2.4 |
| Y169 )( L190 | 7.9 | 9.7 | 8.4 | 16.1 | 19.5 | 20.5 | 4.6 | 4.4 | 2.7 |
| L172 )( A177 | 33.2 | 32.8 | 32.4 | -5.5 | -2.5 | -2.2 | 10.0 | 12.3 | 14.7 |
| T186 )( M214 | 22.0 | 22.9 | 24.4 | -2.7 | -0.7 | -1.1 | 3.4 | 4.1 | 5.4 |
| E187 )( M214 | 24.1 | 21.8 | 21.9 | 0.3 | 0.2 | -0.2 | 2.1 | 1.4 | 0.7 |
| V191 )( M214 | 24.3 | 17.9 | 15.9 | 16.0 | 15.7 | 16.5 | -1.0 | -0.6 | -2.5 |
| C198 )( V221 | 11.2 | 14.9 | 7.3 | 29.9 | 27.7 | 27.3 | 3.3 | 6.0 | 3.2 |
| C218 -H- V221 | 8.6 | 10.4 | 9.0 | 22.2 | 22.7 | 23.1 | 3.1 | 5.9 | 3.6 |
| C218 )( V221 | 5.1 | 8.8 | 1.2 | 18.8 | 20.0 | 20.1 | 1.1 | 4.4 | 1.7 |

**Fig. 9.** Difference in frequency of non-bonded interactions present between secondary structural features of capsid monomers in $HH_{IP6}$, $HH_{PF74}$, and $HP_{apo}$ systems, with respect to monomers in the $HH_{apo}$ system, at low, medium, or high-tilt ranges.

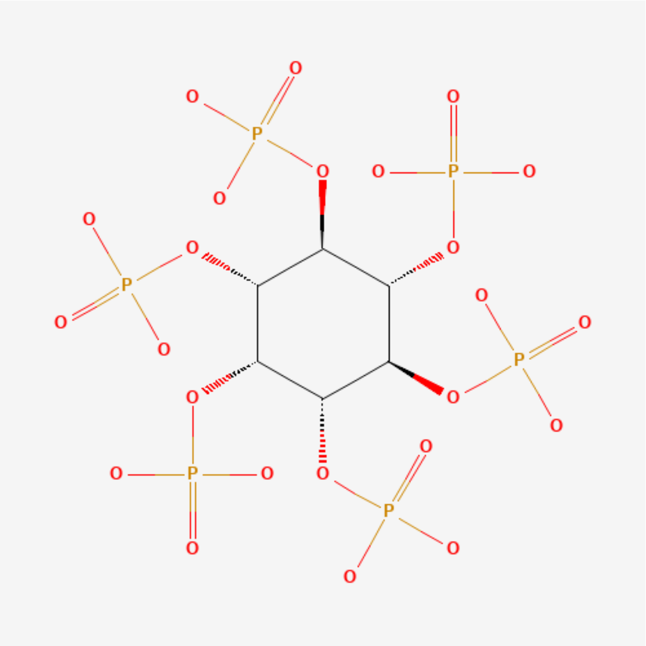


**Fig. 10.** Protonated IP6 (`https://pubchem.ncbi.nlm.nih.gov/compound/21584050#section=2D-Structure`)

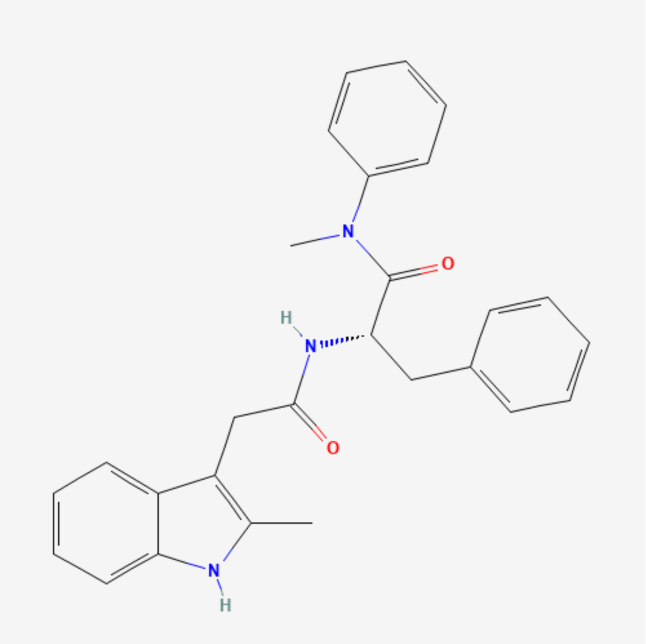


**Fig. 11.** Protonated PF74 (`https://pubchem.ncbi.nlm.nih.gov/compound/49800090#section=2D-Structure`)

### Bibliography


1. Zhao, G.; Perilla, J. R.; Yufenyuy, E. L.; Meng, X.; Chen, B.; Ning, J.; Ahn, J.; Gronenborn, A. M.; Schulten, K.; Aiken, C.; Zhang, P. Mature HIV-1 capsid structure by cryo-electron microscopy and all-atom molecular dynamics. *Nature* **2013**, *497*, 643–6.
2. Dick, R. A.; Zadrozny, K. K.; Xu, C.; Schur, F. K. M.; Lyddon, T. D.; Ricana, C. L.; Wagner, J. M.; Perilla, J. R.; Ganser-Pornillos, B. K.; Johnson, M. C.; Pornillos, O.; Vogt, V. M. Inositol phosphates are assembly co-factors for HIV-1. *Nature* **2018**, *560*, 509–512.
3. Dolinsky, T. J.; Nielsen, J. E.; McCammon, J. A.; Baker, N. A. PDB2PQR: an automated pipeline for the setup of Poisson-Boltzmann electrostatics calculations. *Nucleic Acids Res* **2004**, *32*, W665–7.
